# Exploring the C(2)M Cohesin Complex: Structure, Dynamics, and Ability to Facilitate Assembly of the Synaptonemal Complex

**DOI:** 10.1101/2025.04.21.649846

**Authors:** Margaret Howland, Helen Nguyen, Daria Mitri, Justin Mathew, Nikunj Patel, Mercedes R. Gyuricza, Janet K Jang, Kim S. McKim

**Author notes:** Corresponding author: Kim S. McKim, 190 Frelinghuysen Road, Piscataway, NJ-08854.

## Abstract

There are two meiotic cohesin pathways that regulate synaptonemal complex (SC) assembly in *Drosophila*. We previously proposed that C(2)M, which is required for SC assembly, is the only meiosis-specific component of a complex that includes Stromalin (SA), Nipped-B, SMC1 and SMC3. This model also predicts that specific residues within the C-terminus and N-terminus of C(2)M should interact with SMC1 and SMC3 to form a ring structure that may regulate the ability of C(2)M to facilitate SC assembly. Through mutant analysis, our results show several residues known to interact with SMC1 or SMC3 are critical for SC formation, suggesting that C(2)M may require a ring structure to perform meiosis-specific functions such as the formation of SC. We also show that SA colocalizes with and depends on C(2)M. However, the dynamics of C(2)M differ from SA and the SMCs in a way that suggests C(2)M regulates the dynamics and chromosome loading of the other cohesin proteins SMC1 and SA. Consistent with this conclusion, our results suggest that C(2)M can promote chromosome localization of the other cohesin components, and can induce SC assembly when ectopically expressed in germline mitotic cells.

## Introduction

Homologous chromosomes in meiosis are held together by the synaptonemal complex (SC) (Lake and Hawley 2012). SC assembly is required to stabilize pairing of homologous chromosomes and is required for crossing over. The SC is composed of two lateral elements (LE) joined by a central transverse filaments (TF). The LEs, or axial element, are also at the interface between the SC and the chromatin, sometimes called the chromosome axis. The meiotic chromatin is organized into a linear array of loops, wherein recombination is proposed to occur, that are anchored to the meiotic chromosome axis or axial element (Cahoon and Hawley 2016b; Ito and Shinohara 2022; Zickler and Kleckner 2023).

Cohesin-related proteins are a crucial component of meiotic chromosomes and facilitate assembly of axis and the SC (Ur and Corbett 2021). In mitotic cells, the core cohesin complex includes two subunits of the structural maintenance of chromosomes (SMC) protein family, SMC1 and SMC3 (Losada *et al*. 1998; Losada and Hirano 2005) and a kleisin family protein RAD21/SCC1. Kleisins are a family of eukaryotic and prokaryotic proteins that interact with the SMC proteins through their N- and C-terminal domains to form a ring-like structure (Schleiffer *et al*. 2003). In addition, this core complex interacts with three HAWK (Heat repeat proteins Associated With Kleisin) proteins Scc3/Stromalin, Scc2/Nipped-B and PDS5 (Wells *et al*. 2017; Ishiguro 2019).

In contrast to mitotic cohesin complexes, meiotic cells often have two cohesin complexes (Cahoon and Hawley 2016a). In mammals, these are characterized by meiosis-specific kleisin subunits REC8 (Parisi *et al*. 1999; Eijpe *et al*. 2003; Lee *et al*. 2003) and RAD21L (GutiÉrrez-Caballero *et al*. 2011; Ishiguro *et al*. 2011; Lee and Hirano 2011). In addition, other mitotic cohesin complex subunits may be replaced by meiosis-specific cohesin subunits, although this is not as highly conserved as Kleisin replacement. For example, mouse mitotic cohesin SMC1ɑ and SA1/SA2 are replaced by meiosis-specific cohesin subunits SMC1β (Revenkova *et al*. 2001) and SA3/STAG3 (Bayes *et al*. 2001; Prieto *et al*. 2001). Because the SA and SMC subunits in some organisms are not meiosis specific, the variable kleisin subunits may confer specificity to meiotic cohesin complexes.

The two meiotic cohesin complexes In *Drosophila* contain SMC1 and SMC3, but differ in their meiosis-specific kleisin and SCC3/Stromalin subunits (Tanneti *et al*. 2011). The complex that regulates meiotic sister chromatid cohesion contains the kleisin SOLO, the SA-like protein SUNN, and ORD (Webber *et al*. 2004; Yan and Mckee 2013; Krishnan *et al*. 2014; Gyuricza *et al*. 2016). With its cohesion function, the SUNN-SOLO cohesin complex in *Drosophila* resembles REC8 cohesin complexes in mammals.

The other complex involves the kleisin C(2)M (crossover suppressor on 2 of Manheim). Several lines of evidence suggest that C(2)M, Stromalin/SCC3, and Nipped-B/SCC2, function in one complex with SMC1 and SMC3 (Manheim and Mckim 2003; Heidmann *et al*. 2004) (Gyuricza *et al*. 2016). C(2)M physically interacts with SMC3 (Heidmann *et al*. 2004), and oocytes lacking C(2)M, SA or Nipped-B exhibit similar SC defects, where only patches of the SC forming and persisting throughout prophase. When either C(2)M or Stromalin is mutated, sister centromere cohesion is intact while SC formation and crossovers are impaired (Manheim and Mckim 2003; Gyuricza *et al*. 2016), suggesting that the C(2)M-Stromalin complex is primarily responsible for homologous chromosome interactions rather than sister chromatid cohesion. Electron microscopy shows C(2)M in the chromosome axis (Anderson *et al*. 2005) and colocalizes with the SC protein C(3)G (Manheim and Mckim 2003; Gyuricza *et al*. 2016). C(2)M localization is also reduced in the absence of C(3)G while overexpression of C(2)M results in more SC assembly, suggesting important regulatory interactions between meiotic cohesins and SC assembly (Manheim and Mckim 2003). Thus, C(2)M and RAD21L in mammals may have similar functions (Ishiguro *et al*. 2014).

These results suggest a conserved characteristic of meiosis in a broad range of organisms is two cohesin meiotic cohesin complexes, one for cohesion and the other for organizing the meiotic chromosome axis, and SC assembly (Cahoon and Hawley 2016a; Ur and Corbett 2021). Unlike cohesins for cohesion like Rec8, C(2)M and *C. elegans* Coh-3/ Coh-4 are dynamic during prophase, with subunits being added to and dissociating from the chromosomes throughout pachytene prophase (Gyuricza *et al*. 2016; Castellano-Pozo *et al*. 2023). In this paper, we conducted a mutational analysis to test the hypothesis that C(2)M interacts with SMC1, SMC3 and Stromalin. We also examine the dynamics of these proteins and found that C(2)M has a critical role in regulating the loading of the other cohesin subunits. In fact, expression of C(2)M maybe sufficient to promote assembly of SC in germline cells.

## Results

The oocytes in the germarium are arranged in temporal order, with the earliest stages being more anterior and the later stages are more posterior (Figure 1A). This arrangement allows us to observe a progression of time points, from the premeiotic mitosis (region 1) and premeiotic S-phase through the end of pachytene (Hughes *et al*. 2018). SC assembly is initiated during of meiotic prophase (zygotene), which occurs in region 2a of the *Drosophila* germarium. SC assembly is also completed (pachytene) in region 2a, with the formation of the SC along the length of each chromosome arm, and continues into regions 2b and 3 of the germarium.

**Figure 1:**
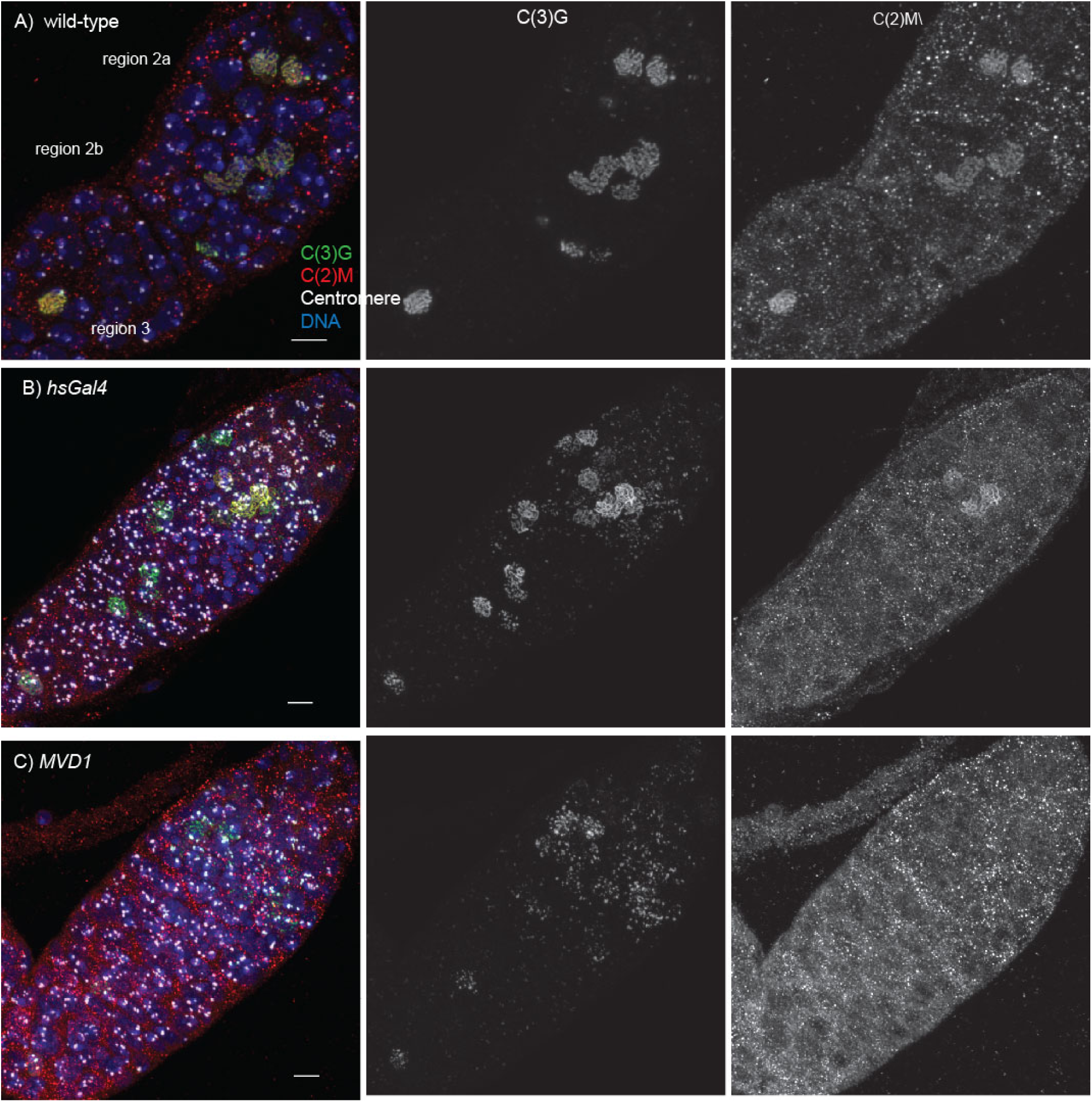
Organization of the germarium and heat shock induced RNAi of C(2)M. A) Wild-type germarium, with C(3)G is in green, C(2)M in red, centromeres (CENP-C), DNA in blue, and the scale bar is 5 µm. Earlier stages (region 2a) are towards the top and later stages (region2b and 3) are towards the bottom. B) Germarium 24 hours after heat shock induced of *c(2)M^RNAi^.* C) Germarium with *c(2)M^RNAi^* under the control of *MVD1*.

### Effect on SC of C(2)M depletion during pachytene

We have shown that C(2)M is dynamic and can be loaded onto meiotic chromosomes throughout pachytene (Gyuricza *et al*. 2016). In these experiments, heat shock induced C(2)M expression was used to observe incorporation after the SC has been assembled. To determine if C(2)M expression must be maintained after the SC has assembled, we used an inducible RNAi system. With an shRNA expressed with *MVD1* to knock down *c(2)M* in all germ cells, a phenotype similar to null alleles was observed, only patches of C(3)G were observed in all stages of meiotic prophase (Figure 1B). For temporal restriction of RNAi, shRNA was expressed using *hsGal4*, with a one hour heat shock followed by dissection 24 hours later. In these ovaries, normal C(2)M and C(3)G was observed in early region 2a (Figure 1C). It is likely that these oocytes entered meiosis after the heat shock (Gyuricza *et al*. 2016). Later in regions 2a and 2b, depletion of C(2)M was observed, but surprisingly, threads of C(3)G remained. Thus, SC structure was maintained even when C(2)M was depleted. Eventually, however, SC structure was effected as in region 3 as shown by the appearance of C(3)G in patches. These results suggest that, while C(3)G localization can persist for a short time following C(2)M depletion, C(2)M must be incorporated in pachytene to maintain complete SC between homologous chromosomes.

### Mutational analysis of C(2)M domains

To test the model that C(2)M interacts with SMC1 and SMC3 in a ring structure, while also conveying meiosis-specific functions in SC assembly, a mutational analysis was conducted. This analysis was based on the conservation of Kleisin proteins, which includes highly conserved amino acid residues in the C(2)M C-terminal and N-terminal domains (Schleiffer *et al*. 2003; Haering *et al*. 2004; Arumugam *et al*. 2006; Gligoris *et al*. 2014) (Figure S 1). Mutants were generated on an a wild type *c(2)M* transgene tagged with HA at its N-terminus (Manheim and Mckim 2003; Gyuricza *et al*. 2016) (Figure 2A). Mutant variants of *c(2)M* were expressed under the control of the UAS system in a wild-type or mutant (*c(2)M^EP^*/*c(2)M^910^*) background. Wild-type HA-C(2)M localized properly to the chromosome axis and rescued the SC assembly and nondisjunction phenotypes of *c(2)M* mutants (Table 1, Figure 2B). These data indicate that the *c(2)M* transgene provides the wild type function of the C(2)M protein.

**Figure 2:**
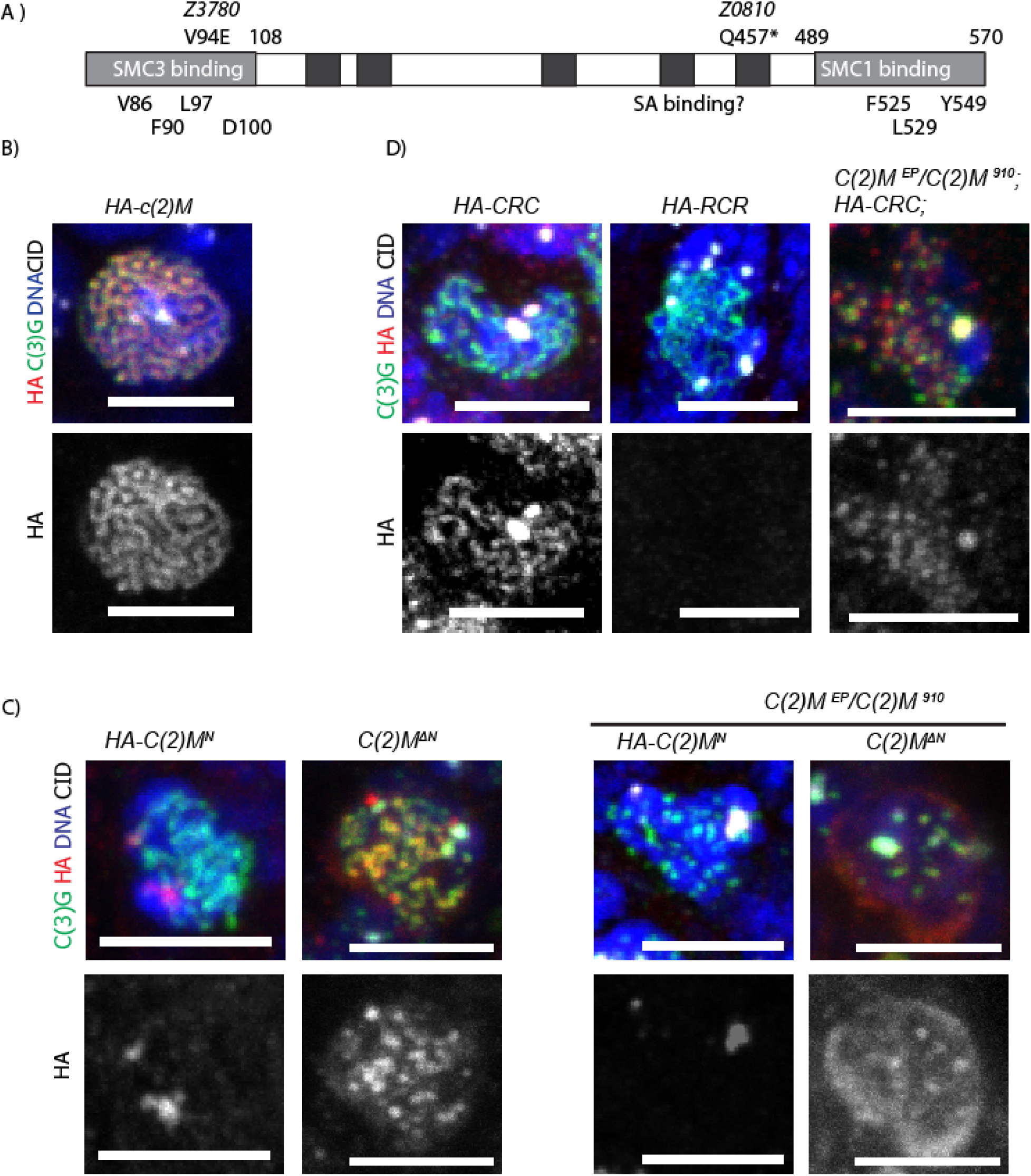
Domain analysis of C(2)M. A) Schematic of C(2)M structure. B) Expression of HA-C(2)M, a wild-type coding region of *c(2)M* fused to 3XHA. C(3)G is in green, C(2)M mutants in red, CID in white, DNA in blue, and the scale bar is is 5 µM. B) HA tagged N terminal domain, or N-terminal domain deletion, of C(2)M in wild-type backgrounds and *c(2)M* mutant backgrounds. D) Expression of VTD/RAD21 – C(2)M fusion proteins. HA-CRC is a fusion with C(2)M N- and C-terminal domains and VTD central domain. HA-RCR is a fusion with C(2)M central domain and VTD N- and C- terminal domains. C(3)G is indicated in green, mutant C(2)M indicated in red, and DNA in blue.

**Table 1.** Nondisjunction in c(2)M mutants in a *c(2)M/c(2)M* mutant background.

|  |  | Oocyte genotype |  |  |  | NDJ (%) |
| --- | --- | --- | --- | --- | --- | --- |
| Genotype |  | XXY ♀ | XO ♂ | XX ♀ | XY ♂ |  |
| N-terminus | HA-c(2)M | 3 | 3 | 998 | 919 | 0.6 |
|  | ΔN | 379 | 361 | 3216 | 2793 | 19.8 |
|  | V86K | 148 | 132 | 1914 | 1605 | 13.7 |
|  | F90A | 1 | 3 | 841 | 786 | 0.5 |
|  | L97K | 142 | 125 | 1685 | 1307 | 15.1 |
|  | D100K | 2 | 1 | 1312 | 1145 | 0.2 |
| c(2)M CRC |  | 39 | 60 | 614 | 484 | 15.3% |
| C-terminus | F525A | 0 | 0 | 461 | 465 | 0.0 |
|  | F525R | 65 | 87 | 582 | 528 | 21.5 |
|  | L529A | 50 | 56 | 985 | 869 | 10.3 |
|  | L529R | 114 | 121 | 1249 | 1001 | 17.3 |
|  | Y549A | 7 | 3 | 1617 | 1333 | 0.7 |
| L97K, L529R |  | 135 | 147 | 1358 | 1132 | 18.5 |

It is not known which domain(s) of C(2)M are required for the meiosis-specific function of SC assembly. In order to address this question, we expressed each of the domains individually with an HA-tag. The N terminal domain of C(2)M (*c(2)M^N^*) localized to the one or two spots in both wild-type and *c(2)m* mutant backgrounds (Figure 2C). Based on the centromere marker CID/ CENP-A, the N terminal domain localizes to the centromeres of the oocyte. These results are surprising because full-length C(2)M localizes specifically to the chromosomes arms and not the centromeres. Ectopic localization of the C(2)M^N^ did not exhibit a dominant phenotype (Table S 1). These results suggest that the N terminal domain may promote interaction with the SMC3, but other domains take part in directing C(2)M to its proper locations as well as facilitating the assembly of the SC.

Expression of the central and C-terminal domains did not result in localization to the chromosomes (Figure S 2). The central SA-binding domain of C(2)M did not localize to the chromosomes. While foci of staining were observed, they did not correspond to any chromosomal location. This data indicates that the SA-binding domain alone cannot localize to chromosomes, but may form aggregates within the oocytes. Expression of the C terminal domain of C(2)M did not result in any chromosome localization. This was observed across multiple lines of the transgenes, suggesting the C-terminal domain protein fragment was unstable, and could indicate the C terminal fragment is rapidly degraded.

Because the C-terminal domain was unstable, we generated a deletion of the C(2)M N-terminal domain, *c(2)M^ΔN^*, which is predicted to abolish C(2)M interactions with SMC3. In a *c(2)M* mutant background, both the cytology (Figure 2C) and nondisjunction levels of *c(2)M^ΔN^* (Table 1) were comparable to that of *c(2)M* null alleles. SC formation in *c(2)M^ΔN^* oocytes was interrupted, with C(3)G forming in foci instead of threads, similar to null alleles. Expression of *c(2)M^ΔN^* in wild-type background did not result in a dominant phenotype (Table S 1), and C(3)G localized properly in threads. The C(2)M^ΔN^ protein localized in foci on the chromosomes and was more abundant on the chromosomes in a wild-type C(2)M background compared to the mutant background. Thus, in the absence of the N-terminal domain, there was a defect in cohesin loading and SC assembly. The increased localization in a wild-type background suggests the C(2)M^ΔN^ mutant protein can interact with intact cohesin complexes, or the SC itself.

### Fusions of C(2)M and Rad21/Vtd

The Kleisin central domain is believed to interact with regulatory subunits such as Stromalin (Higashi and Uhlmann 2022). To investigate if the meiosis-specific functions of C(2)M in SC assembly are within the N- or C-terminal domains that interact with the SMC proteins or within the central domain, we constructed fusions of *c(2)M* and *Rad21* (= *vtd,* the mitotic kleisin protein in *Drosophila*). Two versions of the fusions were made; one with the N and C termini of C(2)M and the Rad21 central region (CRC), and one with the N and C termini of Rad21 and the SA-Binding domain of C(2)M (RCR). The RCR protein was not detected by immunofluorescence, suggesting it is unstable and degraded.

The CRC protein localized in threads to the chromosomes in a C(2)M wild type background (Figure 2D). Surprisingly, CRC localized to the centromeres as well as the chromosome arms, in contrast to C(2)M, which only localizes to the chromosomes arms. However, CRC did not rescue the SC assembly defects in a *c(2)M* mutant background. Instead, the CRC protein localized in patches that co-localized with C(3)G on the chromosome arms as well as the centromeres. Thus, the CRC protein can localize to meiotic chromosome axis but does not promote SC assembly. Consistent with these results, CRC mutant females had high levels of nondisjunction compared to wild type flies (15.3%, Table 1) but did not have dominant effects (Table S 1). These results suggest that the central region of C(2)M is required for the meiotic localization pattern and the SC assembly function of C(2)M.

To determine if the CRC fusion could function in mitotic cells like VTD, we expressed it in somatic cells using a ubiquitous expressing Gal4 (*Tub-Gal4*). While the original goal of this experiment was to determine if expression of CRC could rescue the lethality of *vtd/rad21* RNAi flies, we found that ubiquitous expression of CRC caused lethality (Table 2). This suggested that the CRC protein interacts with the SMC proteins in mitotic cells but interferes with normal cohesion regulation.

**Table 2:** C(2)M Rad21 Fusion Mitotic Viability Test.

| Genotype | Sb | Sb+ | Total Progeny |
| --- | --- | --- | --- |
| *CRC86* | 1816 | 0 | 1816 |
| *CRC3* | 1045 | 0 | 1045 |
| *HA-C(2)M* | 317 | 283 | 600 |

CRC mutant is a fusion of Rad21/VTD central domain with C(2)M terminal domains. In each experiment, the wild-type or mutant transgene males were crossed to *TubGal4/TM3, Sb* females. The Sb+ progeny express the *C(2)M* transgene in all somatic cells. The absence of Sb+ progeny indicates this genotype is viable.

### Critical residues in C(2)M for synaptonemal complex assembly

To test the hypothesis that a C(2)M cohesin complex ring structure is important for SC assembly, we examined C(2)M variants with mutations in residues predicted to disrupt interactions between C(2)M and one of the SMC proteins. While the central region of C(2)M is poorly conserved, residues in the N- and C-terminal domains are highly conserved across all kleisin proteins (Schleiffer *et al*. 2003) (Haering *et al*. 2004; Arumugam *et al*. 2006; Gligoris *et al*. 2014) (Figure S 1). The N-terminal domain of Scc1 folds into three alpha-helices, with a 34-residue α3 (R69-M102) forming a long helical bundle with SMC3’s coiled-coil domain found in its globular head domain (Gligoris *et al*. 2014). The face of Scc1’s α3 that contacts SMC3’s coiled-coil region is conserved (Schleiffer *et al*. 2003). When conserved residues (corresponding to *C(2)M* F90 and L97) in this region were substituted with lysine, it caused lethality and reduced the association between Scc1 and SMC3 (Gligoris *et al*. 2014). The C-terminus of Scc1 is a winged-helix domain (WHD) and hydrophobic residues like F528 and L532 make contact with the two beta-strands of SMC1’s globular head (Haering *et al*. 2004) (Arumugam *et al*. 2006). Mutants of these conserved residues were examined in a wild-type or C(2)M mutant background for SC assembly and the percentage of homologous chromosome nondisjunction (Table 1). In addition to the amino acids noted above, we also mutated three other resides *V86, D100* and *Y548* that are conserved but not previously associated with SMC1/3 interactions.

In a *c(2)M* mutant background, the *F90A*, *D100K* (Figure 3), *F525A*, and *Y549A* (Figure 4) mutants showed wild-type-like SC assembly. C(2)M colocalized with C(3)G in threads along the chromosome arms. Furthermore, *F90A*, *D100K*, *F525A*, and *Y548A* in a *c(2)M* null mutant background showed levels of nondisjunction similar to wild-type (Table 1).

**Figure 3:**
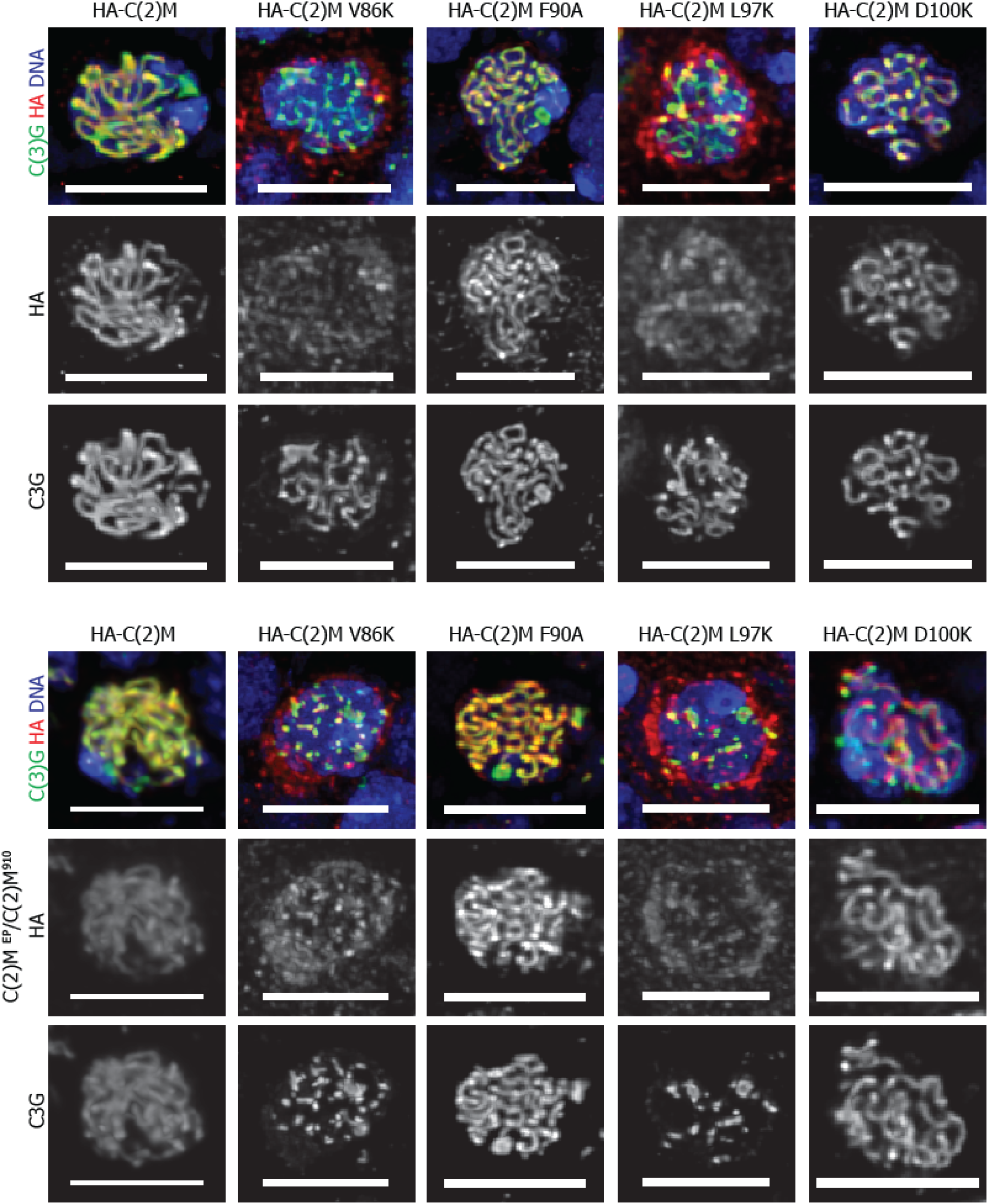
Mutants in the N-terminal domains of C(2)M. A) Expression of wild-type or *HA-c(2)M* mutants in the N terminal domain in a wild-type *c(2)M* background. B) Expression of wild-type or *c(2)M* mutants tagged with HA at the N terminal domain in a *c(2)M* mutant background. C(3)G is in green, HA / C(2)M is in red, DNA in blue, and the scale bar is 5 µM.

**Figure 4:**
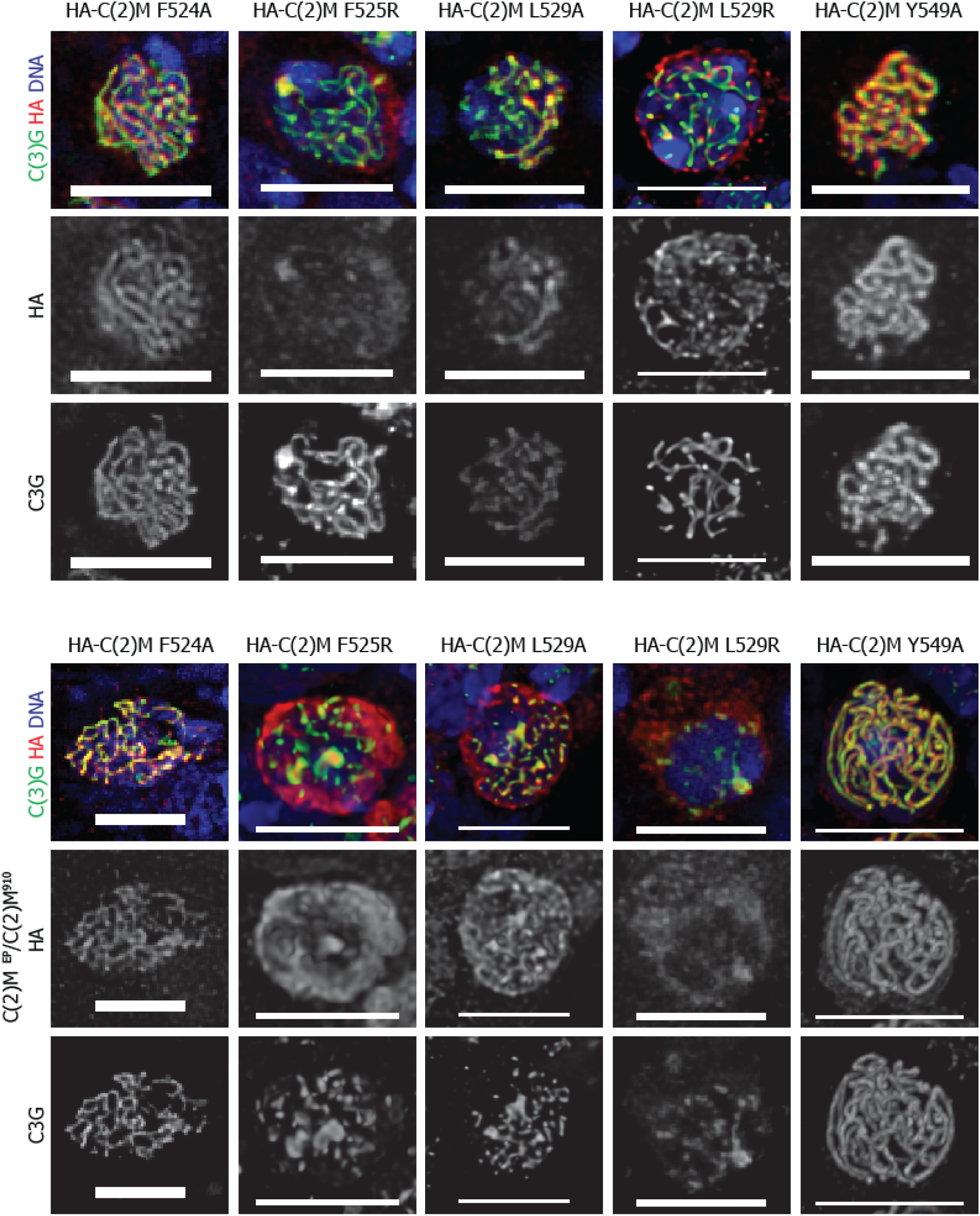
Mutants in the C-terminal domains of C(2)M. A) Expression of *HA-c(2)M* mutants in the C-terminal domain in a wild-type *c(2)M* background. B) Expression of *c(2)M* mutants in the C-terminal domain in a *c(2)M* mutant background. C(3)G is in green, HA / C(2)M is in red, DNA in blue, and the scale bar is 5 µM.

In contrast, *c(2)M* point mutations *V86K* and *L97K* in the N-terminus and *F525R*, *L529A*, and *L529R* on the C-terminus had only patches of C(3)G throughout regions 2 and 3 of the germarium, indicating defects in SC assembly. Consistent with the defects in C(3)G, C(2)M localization was also abnormal. Rather than localizing in threads along the chromosomes, the mutant protein was present in only a small number of nuclear foci. These defects were associated with a significant amount of mutant cytoplasmic protein. This was not observed when C(2)M localized in threads, such as the wild-type C(2)M protein and in the *F90A*, *D100K*, *F525A*, and *Y548A* mutants. Thus, the failure to localize was associated with increased accumulation in the cytoplasm. We also observed differences between N- and C-terminal mutants. The N-terminal *V86K* and *L97K* mutants had stronger localization to foci on the chromosomes. The C-terminal mutants *F525R* and *L529R* had stronger localization to the centromeres, similar to the *c(2)M^N^* mutant (Figure S 3).

The point mutants that showed abnormal SC assembly (*V86K*, *L97K*, *F525R*, *L529A*, and *L529R*) also had high levels of nondisjunction in mutant *c(2)M* background (Table 1). The fact that *F525R* and *L529R* mutants had a more severe defects than *F525A* and *L529A* indicates the charge of the amino acid is important. In contrast, nondisjunction levels of all *c(2)M* point mutants in a wild-type background was similar to the wild-type control (Table S 1). Similarly, C(3)G localization was not effected by expression of the mutant C(2)M proteins in a wild-type background (Figure 3, Figure 4). Thus, the *c(2)M* mutants did not have a dominant phenotypes.

In the analysis of the C(2)M domains above, a fragment containing only the central domain failed to localized to the chromosomes. To address the function of the central domain, we generated a double mutant with a mutation in each SMC binding domain, *L97K* and *L529R*. The double mutant *L97K, L529R* had nondisjunction (18.42%) and cytological defects that were similar to the *c(2)M* point mutations in each domain (Table 1). Notably, the localization of this mutant protein was similar to the single mutants, including the absence of centromere localization (Figure S 2, Figure S 3). Given the two SMC binding domains were mutated, the foci of chromosome localization could be due to the central domain of C(2)M.

In summary, these results suggest that V86, L97, F525, L529 are critical residues in C(2)M that are important for facilitating SC assembly, possibly through mediating interactions with SMC1 and SMC3 and forming a cohesin ring structure. In contrast, and despite being conserved, F90, D100, and Y548 may not be as important for these interactions.

### SA Localization is Dependent on C(2)M and Nipped-B but not VTD

In addition to interactions with SCM1/3, we previously proposed that Stromalin is required for C(2)M cohesin function. To characterize SA localization during meiosis, we created a MYC-tagged transgene under the control of *UASp*. SA-MYC in a wild-type background expressed with MVD1, a germline-specific promoter, was detected in a thread like pattern colocalizing with C(3)G (Figure 5A). SA was also found in a more diffuse cytoplasmic pattern in several other cells. The diffuse SA-MYC staining could be caused by excess transgene expression in the nucleus. When SA-MYC was expressed with MVD1 in a *c(2)M* mutant background, all the SA-MYC localization was abolished, even in regions of diffuse staining (Figure 5B). These data indicate that SA localization is dependent upon C(2)M. The absence of diffuse or cytoplasmic SA-MYC staining could indicate that SA stability depends on interacting with C(2)M, even if not localized to the chromosomes and in the cytoplasm.

**Figure 5.**
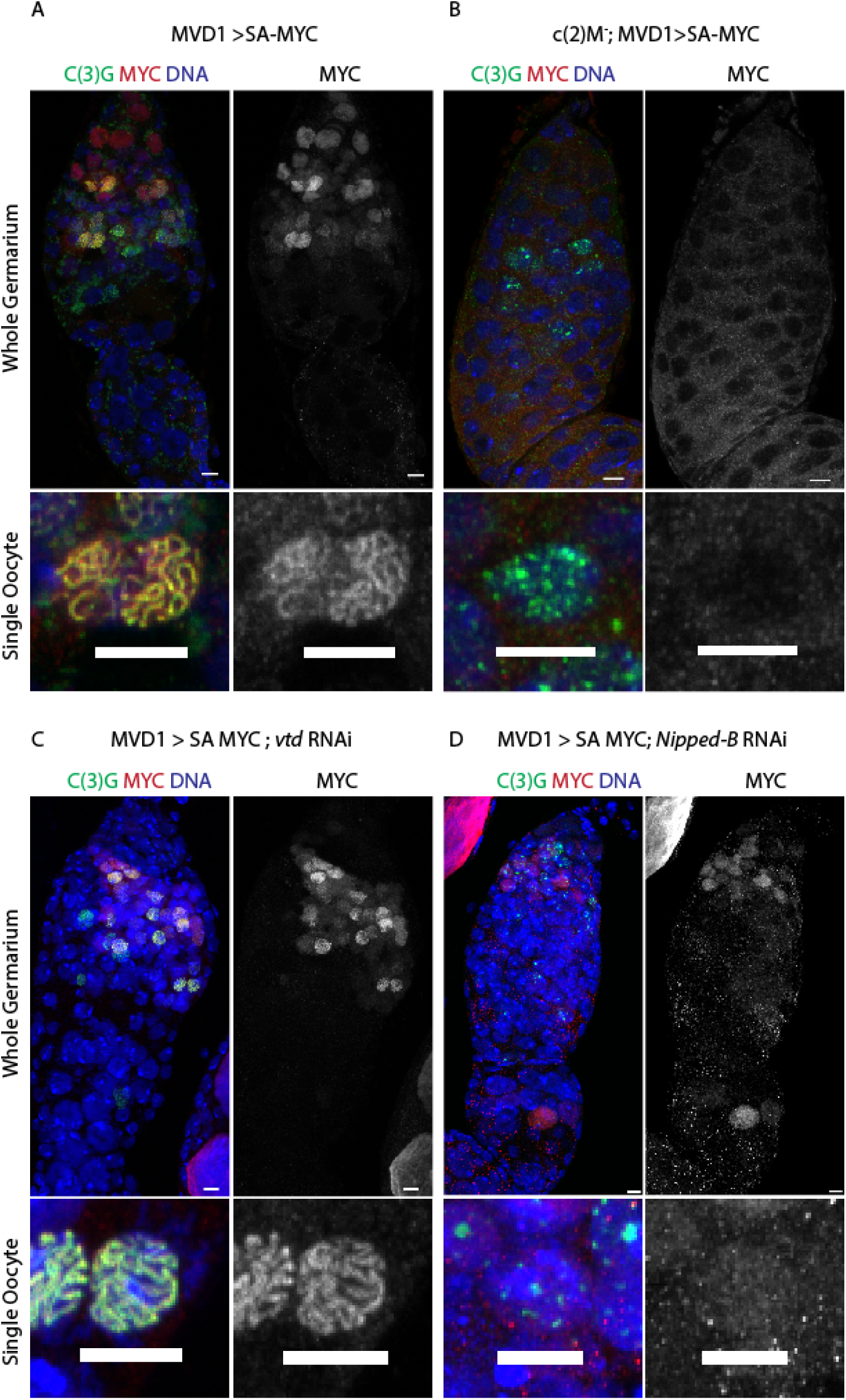
SA-MYC colocalization with C(3)G is dependent on C(2)M and Nipped-B but Not VTD. In all images, C(3)G is in green, SA-MYC is in red, and DNA is in blue. For each genotype, a representative whole germarium is shown above a single oocyte. The genotypes were (A) wild-type, (B) *c(2)M* (C) *vtd*^RNAi^, and (D) *Nipped-B*^RNAi^. Scale bars is 5 µm.

In mitotic cells, SA interacts with the kleisin / RAD21 orthologue VTD. We hypothesized that VTD would not interact with SA during meiotic divisions because *vtd* RNAi has no SC assembly defect (Gyuricza *et al*. 2016). Indeed, we observed that in *vtd* RNAi oocytes, SA localized to the chromosome axis, indicating that VTD is not required for SA localization in meiosis (Figure 5C).

Nipped-B is proposed to regulate the mitotic cohesin complex SMC1/SMC3/VTD/SA. Nipped-B knockdown has a similar cytological phenotype to both SA knockdown and C(2)M mutants (Gyuricza *et al*. 2016), suggesting the meiotic SMC1/SMC3/C(2)M/SA complex interacts with Nipped-B. To test if SA localization is dependent upon Nipped-B, we co-expressed *Nipped-B* shRNA and SA-MYC. In the presence of the *Nipped-B* shRNA, SA-MYC did not localize to the chromosomes (Figure 5D). This result is not as severe as the abolition of SA localization in a *c(2)M* null mutant, which could be because the mRNA message is only depleted by 70% in the *Nipped-B* shRNA (Gyuricza *et al*. 2016).

### SA is required for meiotic crossing over and chromosome segregation

Synapsis defects such as those observed in *SA* RNAi oocytes should result in reductions in crossing over and a corresponding increase in nondisjunction (Lake and Hawley 2012; Gyuricza *et al*. 2016). SA is an essential gene, however, and a strong germline reduction of SA by RNAi results in sterility. To genetically examine chromosome segregation, we used *NGTA* to expresses GAL4 in the germarium but not in later stages (see Methods). Expressing shRNA with *NGTA* was predicted to knock down SA during pachytene, but allow protein expression in late stage oocytes to support embryonic development and fertility. Indeed, high levels of X-chromosome nondisjunction were observed when *NGTA* was used to express *SMC3* or *SA* shRNA in the germarium (Table 3), consistent with a defect in SC assembly and crossover formation.

**Table 3:** Nondisjunction with different GAL4 lines with various shRNA lines.

| RNAi | Oocyte genotype |  |  |  | Nondisjunction |
| --- | --- | --- | --- | --- | --- |
| Genotype | XXY | XO | XX | XY | (%) |
| *w1118* | 2 | 3 | 1855 | 1770 | 0.3 |
| *SMC3 /NGT* | 109 | 122 | 958 | 560 | 23.3 |
| *SA /NGT* | 159 | 153 | 1607 | 773 | 20.7 |
| *c(2)M/ NGT* | 95 | 99 | 992 | 734 | 15.9 |
| *c(2)M/ MVD1* | 36 | 38 | 468 | 361 | 15.2 |
| *vtd /NGT* | 1 | 1 | 1101 | 789 | 0.2 |

To measure crossing over, *SA* and *SMC3* shRNA were expressed in females heterozygous for a marked X-chromosome (*y w cv m f • y^+^/ y w*) (Table 4). In these females, one of the X-chromosome centromeres was marked with the *y^+^* marker, which allows for the detection of sister chromatid nondisjunction events. As expected from the nondisjunction data, the RNAi knockdowns of *SA* and *SMC3* had severe reductions in meiotic crossing over (Table 4). Thus, like C(2)M, SA and SMC3 are required for meiotic crossing over. It is notable that nondisjunction of sister chromatids was not detected. This is consistent with SA because it is not predicted to be part of the meiotic complex require for cohesion. However, the lack of sister chromatid nondisjunction in *SMC3* RNAi oocytes may indicate there is a different threshold for cohesion and SC assembly functions. It is possible that a smaller amount of cohesin complexes are sufficient for cohesion than SC assembly function. Alternatively, SMC3 protein required for cohesion may be loaded earlier (Gyuricza *et al*. 2016) or after (Weng *et al*. 2014) when *NGTA* expression effects mRNA levels.

**Table 4:** Crossing over and sister chromatid segregation in *SMC3*, *SA* and *c(2)M* RNAi females.

| Genotype | *cv m*<br>(*mu*) | *m f*<br>(*m.u.*) | Total<br>males | *y* ♀ | NDJ % | Total<br>progeny |
| --- | --- | --- | --- | --- | --- | --- |
| *wild-type* | 21 | 19 | 719 |  | 0.4 | 1614 |
| *SMC3 / NGTA* | 1.2 | 1.6 | 428 | 1 | 35.7 | 1198 |
| *SMC3 / bam* | 6.0 | 4.0 | 201 |  | 26.2 | 478 |
| *wild-type* | 17.8 | 17.6 | 653 |  | 0 | 1464 |
| *SA / NGTA* | 0.22 | 2.0 | 451 |  | 31.4 | 1434 |
| *SA / bam* | 3.2 | 6.8 | 761 |  | 9.47 | 1690 |
| *wild-type* | 27.2 | 20.7 | 558 |  | 3.5 | 1170 |
| *c(2)M / MVD1* | 15.2 | 4.5 | 466 |  | 25.7 | 1399 |
| *c(2)M / NGTA* | 15.2 | 11.0 | 190 |  | 20.1 | 624 |

### SA and SMC1 are dynamic in meiotic prophase

We previously showed that C(2)M can be incorporated into the chromosome axis throughout meiotic prophase (regions 2 and 3) (Gyuricza *et al*. 2016). In contrast, SMC1 only incorporated in late (region 3) prophase oocytes. We performed experiments to determine if the dynamics of SA and C(3)G are similar to SMC1 or C(2)M. Heat shock induced expression was accomplished by crossing *UASP* transgenes to *hsGal4,* the resulting females received a one hour heat shock, and they were dissected 24 hours later.

As observed previously, when *SMC1* expression was induced by heat shock, SMC1 was mostly absent from the chromosomes in region 2a/b oocytes, and appeared in region 3 (Figure 6A, Figure S 4). When SA expression was induced by 1 hour heat shock, a similar staining pattern was observed. Threadlike SA localization was mostly observed in region 3, while non-specific localization appeared throughout the cell cytoplasm in region 2a/2b (Figure 6B, Figure S 4). Similar observations were observed with C(3)G. Following heat shock induced expression of *c(3)*G, localization to the SC was observed mostly in region 3 oocytes (Figure S 5). These results suggest that cohesin and SC components are most dynamic towards the end (region 3) of pachytene.

**Figure 6:**
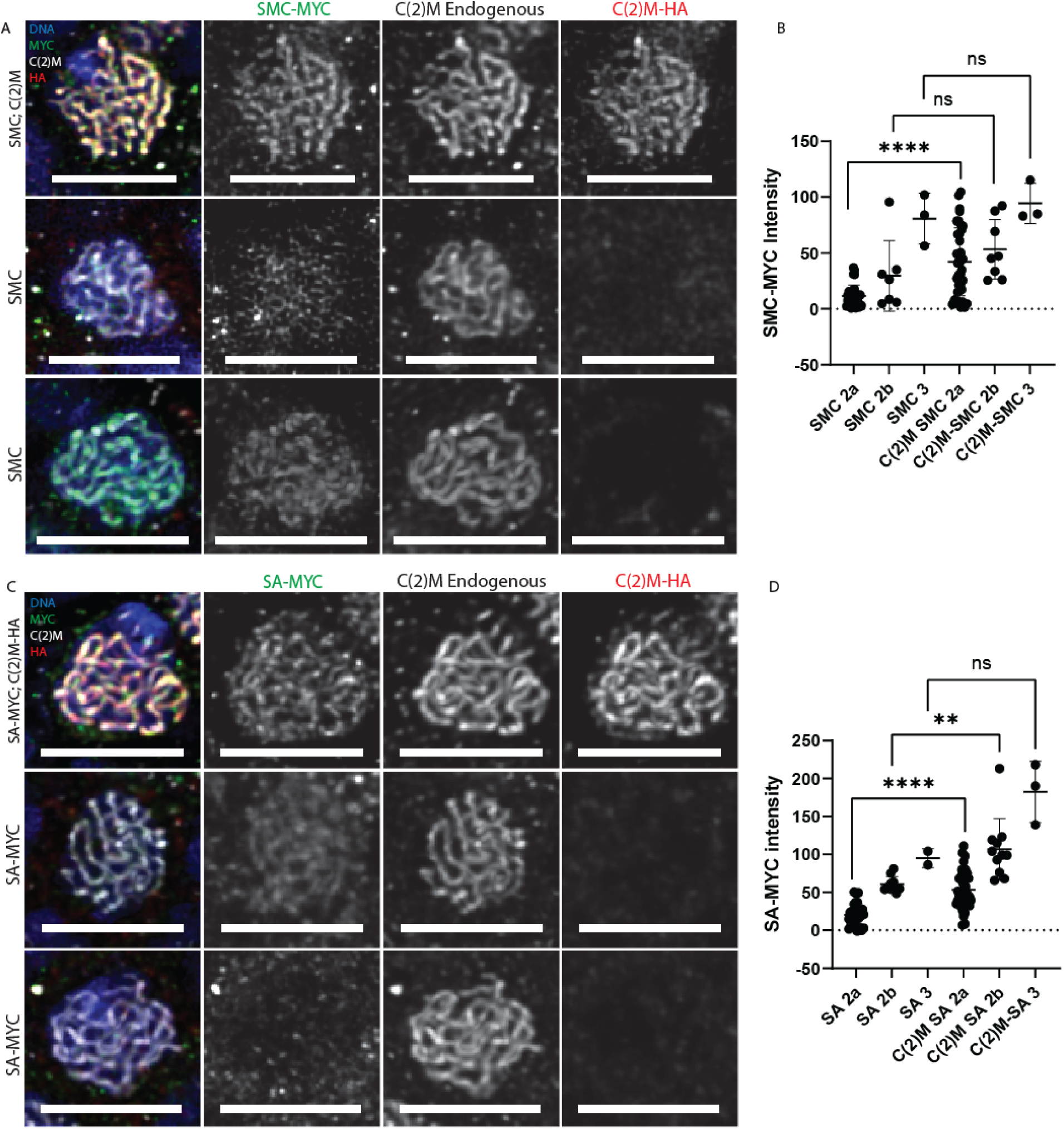
Heat shock induced overexpression of SA and SMC1. A) Colocalization of SMC1-MYC with C(2)M when overexpressed with C(2)M. Cells were stained with SMC3-MYC (green), C(2)M endogenous (white), C(2)M-HA (red), and DNA (blue). B) Quantified SMC1 intensity for each genotype. C) Colocalization of SA-MYC with C(2)M when overexpressed with C(2)M. Cells were stained with SA-MYC (green), C(2)M endogenous (white), C(2)M-HA (red), and DNA (blue). D) Quantified SA intensity for each genotype.

These observations are different than with C(2)M, which appears in most region 2a and 2b oocytes following heat shock (Gyuricza *et al*. 2016). Surprisingly, when *SA* or *SMC1* transgenes were co-expressed with *c(2)M*, there was an increase in SA or SMC1 localization in a threadlike pattern in regions 2a and 2b oocytes (Figure 6A-C). These results suggest that the limiting factor for cohesin localization in region 2a or 2b is C(2)M, and the dynamics of SA or SMC1 on meiotic prophase chromosomes depends on C(2)M.

### Inducing germ-line cells to form SC

The heat shock data suggests that C(2)M may be a limiting factor in cohesin loading and SC assembly. Furthermore, while C(2)M and C(3)G normally localize in threads beginning germarium in region 2a, we have consistently observed threads of C(2)M and C(3)G in more cells per cyst than normal when C(2)M is overexpressed under control of MVD1 (Manheim and Mckim 2003). These results suggest that the expression of C(2)M may be sufficient to initiate SC assembly. In normal ovaries, *c(2)M* is only transcribed in region 2a or later (Manheim and Mckim 2003). Region 1 cells that divide mitotically have cohesin complexes that contain SMC1/SMC3/SOLO/SUNN and SMC1/SMC3/RAD21/SA (Gyuricza *et al*. 2016). However, when a C(2)M-HA transgene is expressed using MVD1, C(2)M is incorporated along the chromosome in region 1 germ cells (Figure 7A).

**Figure 7:**
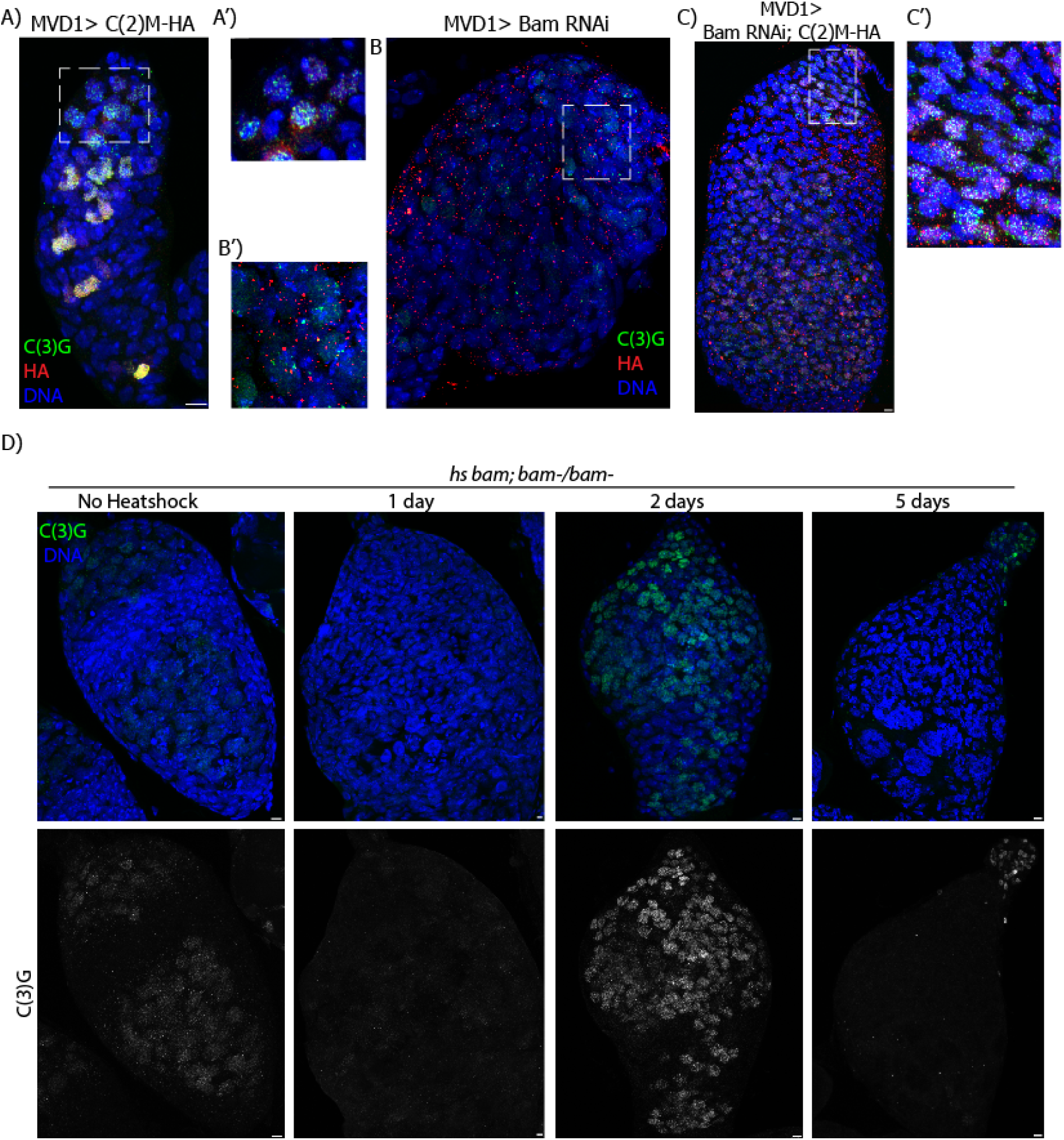
Expression of C(2)M in meiotic germline induced SC assembly. C(3)G is in green and marks the SC. HA is in red and DNA is in blue. A shows C(2)M-HA expression in a wildtype background using the MVD1 promoter, which begins expression in the pre-meiotic divisions. A’ is a blow up of the pre-meiotic region, region 1, and shows SC assembly in this region. B shows the *bam* RNAi also expressed with MVD1. In this panel there are no cells forming much SC and the germarium is much larger than normal. B’ shows a blow up of the earliest region in the *bam* RNAi. C shows C(2)M-HA expressed in a *bam* RNAi background. SC is able to assemble in this case as well. Scale bars are 5µm.

To test if C(2)M expression can drive SC assembly, we used an shRNAi against *bag of marbles* (*Bam*). In the absence of *bam,* gametogenesis is not completed, oocytes are not made, and instead, a large amount of mitotically dividing germline cells proliferate (Mckearin and Spradling 1990). These cells do not show C(3)G staining, indicating that the cells are not entering meiosis (Figure 7B). Interestingly, when the C(2)M-HA transgene was expressed with MVD1 in a *bam* shRNAi background, most of the mitotic cells in the germline started to localize C(2)M and C(3)G on the chromosomes, suggesting C(2)M is sufficient to initiate SC assembly in the mitotic germline (Figure 7C).

These results suggest that the *bam* mutant cells are capable of some meiotic development. Therefore, we investigated if it was possible to induce a large amount of cells in the germarium to enter meiosis at the same time. We induced expression of a *bam* transgene with heatshock in a *bam* mutant background (Ohlstein and Mckearin 1997). Females were heatshocked for one hour pulse and then the ovaries were fixed 1,2 or 5 days later. The *bam* mutant normally has centromeric patches of C(3)G staining and the germarium are much larger than normal because of an abnormally large amount of premeiotic germ cells. One day after heat shock the *bam* mutant germariums looked the same. However, two days after heat shock there was a large amount of C(3)G that almost formed threads, although it still looked abnormal compared to wild type threads of C(3)G (Figure 7D). At five days after heat shock, the C(3)G staining reverted back to the mutant phenotype. These results suggest that the *bam* mutant germ cells have the capacity to enter the meiotic program.

## Discussion

Most organisms have meiotic cohesins. We undertook this study to understand how meiotic cohesins, such as Drosophila C(2)M, promote SC assembly. C(2)M is the only meiosis specific component of the proposed SMC1/SMC3/C(2)M/SA/Nipped-B complex. Thus, it is likely that C(2)M contains domain(s) that promote SC assembly. We have investigated if the domains of C(2)M required for cohesin ring formation are required for SC assembly. Forming a ring is probably also important for meiosis, which may be required for interactions with chromatin. One feature C(2)M is dynamics during prophase, rather than loading specifically at S-phase. C(2)M appears to have a prominent role in regulating cohesin dynamics and localization of the other complex components.

It is unclear if C(2)M interacts directly with SC proteins. Depleting C(2)M by heat shock induced RNAi showed that C(2)M was lost from chromosomes more rapidly than C(3)G, suggesting C(2)M may be required for loading but not maintenance of the SC. However, we observed multiple examples where C(2)M mutant protein localization was higher in a wild-type with full SC than a *c(2)M* mutant background with patches of SC. This could indicate a direct interaction where SC stabilizes C(2)M, which is also consistent with prior observation that C(2)M localization is reduced in *c(3)G* mutant oocytes (Manheim and Mckim 2003). It is also possible, however, that C(2)M forms multimers, and the wild-type can stabilize localization of mutant proteins.

### Insights into the interaction between C(2)M and SMC1 or SMC3

In the case of SCC1, it has been suggested that an essential first step in cohesin ring formation is binding of the C-terminal domain to SMC1, which is followed by N-terminal domain binding to SMC3 (Arumugam *et al*. 2003). This could explain why the C-terminal domain mutants tended to have a weaker localization to the chromosomes than the N-terminal domain mutants. However, a fragment containing only the C(2)M N-terminal domain was found to localize to the centromeres in oocytes, which is most likely by interacting with SMC3. In addition, mutants with a C-terminal domain mutation had significant centromere localization but were defective in interacting with other chromosomal sites, suggesting the N-terminal domain can bind to the SMC proteins independent of other domains. N-terminal domain mutations, or mutants lacking the N-terminal domain, localized to foci but not strongly at the centromeres. One interpretation of these results is that the C-terminal domain interaction with SMC1 has a role in determining the localization pattern of C(2)M, specifically along the euchromatic arms but not at the centromeres.

We were unable to test the localization of the C-terminal domain on its own because it did not produce stable protein. One possibility is that this fragment may lack nuclear localization signals. C(2)M has predicted NLS sequences at positions 55, 131, 147, and 324. Indeed, a fragment containing the C-terminal domain and the central region, but lacking the N-terminal domain, localized to several euchromatic sites. The central domain could also promote localization to euchromatic sites, and prevent centromere localization. To test the role of the central domain, we made the *L97K,L529R* mutant. The localization resembled the single *c(2)M* point mutations and the level of nondisjunction was not significantly worse than in single *c(2)M* point mutations. Thus, a total loss of SMC interaction may not prevent C(2)M interaction with the chromosomes. An intriguing possibility is that the central domain facilitates interactions with other SC or chromosomal proteins.

Through cytological and genetic nondisjunction assays, we found key residues (V86, L97, F525, and L529) required for C(2)M chromosome localization, SC assembly and resulting crossing over and chromosome segregation. The defects in *V86*, *L97*, *F525*, and *L529* mutants were comparable to the nondisjunction levels *c(2)M* null mutations and to *c(2)MΔN*. As in previous studies, the most effective mutations were replacement with a charged residue rather than Alanine (Haering *et al*. 2004; Arumugam *et al*. 2006; Gligoris *et al*. 2014). Most of these residues are conserved in all Kleisins. An exception is the residue corresponding to V86 is not conserved C. elegans Coh-3. Overall, our findings are consistent with a cohesin ring structure facilitating synaptonemal complex (SC) assembly.

Other amino acid positions tested (F90, D100, Y549) did not have severe meiotic defects and may not be required for interactions with SMC1 or SMC3. These residues are less critical for meiotic function based on the observation that each mutant had mild or no defects in C(2)M localization, SC assembly, or chromosome segregation. While F90 is in a conserved position, it is usually a Y in other Kleisins and is not conserved in *C. elegans*. However, this result was surprising because yeast Y82 is proposed to make an important interaction with SMC3. D100 is conserved in all Kleisins, including *C. elegans*, but had a mild meiotic phenotype. It may not be surprising that the Y548 mutation had a mild phenotype, because this residue does not precisely align with a conserved Y in other Kleisins.

### Meiotic Specificity of C(2)M and toxicity to Mitotic Cells

To test the role of each C(2)M domain in meiosis, we did substitutions with *vtd/Rad21* central and flanking domains. Expression of the RCR construct, which contains the C(2)M central domains but the VTD/RAD21 terminal domains, was not detected on chromosomes. This could reflect a mechanism to exclude the mitotic cohesins from the meiotic chromosomes.

In contrast, the CRC construct, which contains the C(2)M terminal domains but the VTD/RAD21 central domain, localized along the meiotic chromosomes, but did not function in SC assembly. These results suggests that the central domain of C(2)M has a role in regulating localization to the chromosomes and formation of the SC. This could occur through interactions with SA, as it has been shown that a complex involving SA and a region within the Kleisin central domain may initiate interactions with the DNA (Li *et al*. 2018; Higashi *et al*. 2020). Further studies are necessary to determine if a similar region resides in C(2)M. The central region is the most diverged of C(2)M, which is surprising because it probably interacts with SA. SA may not confer any meiosis-specificity, given that SA also has a role in mitotic cohesion. This would also explain why SA homologs in different species interact with different meiotic cohesins. For example, rice SCC3 interacts with Rec8 in Rice (Zhao *et al*. 2024). Interestingly, mammal genomes encode a meiosis-specific SA3 subunit (Prieto *et al*. 2001; Winters *et al*. 2014).

Expression of the CRC construct in somatic cells caused dominant lethality. The reason for this is not clear, especially if the terminal domains of C(2)M can interact with the SMC proteins but the central domains confers meiotic or mitotic specificity. Instead, these results suggest the N-terminal domains have more function than simply interacting with SMC proteins, but also confer meiosis-specific functions. Conversely, expression of mutant forms of C(2)M did not confer dominant meiotic defects, even though these mutant forms were clearly incorporated into the meiotic chromosome axis.

### C(2)M is Rate Limiting for SA and SMC Loading and SC assembly

We previously showed that Stromalin and Nipped-B are required for C(2)M localization (Gyuricza *et al*. 2016). Whereas SMC1/3, SA and Nipped-B are required for mitotic cohesins, we proposed that C(2)M/SA/Nipped-B defines one of two the meiosis specific cohesin complexes. Consistent with this hypothesis, Stromalin localization is similar to and depends on C(2)M and Nipped-B. Furthermore, *SA* and *Nipped-B* depleted oocytes also have similar defects in meiotic recombination and chromosome segregation to *c(2)M*.

We previously observed that the dynamics of SMC3 and C(2)M are different (Gyuricza *et al*. 2016). Heat shock induced expression of C(2)M showed incorporation throughout pachytene, while SMC3 protein was incorporated only in late pachytene (mostly region 3). Here we have shown that the dynamics of SMC3 and SA are dependent on C(2)M. Increased SMC3 and SA localization was observed when coexpressed with C(2)M. Thus, it is likely that SMC3 and SA loading is dependent on C(2)M. Furthermore, one explanation for the cytoplasmic localization observed with overexpression of SMC and SA without C(2)M could be that C(2)M facilitates transport of the other cohesins into the nucleus. This is similar to the proposal the LinE proteins, a component of the meiotic chromosome axis in *S. pombe*, enter the nucleus as one complex (Davis *et al*. 2008; Wintrebert *et al*. 2021). The LinE proteins include Rec25, Rec27, Mug20 and Rec10, but the only NLS is found in Rec10.

These results show that C(2)M regulates the dynamics of other cohesin components and the SC. The analysis of the other cohesins or SC component C(3)G suggest the dynamics of most components of the SC are relatively stable during early pachytene, and increase later pachytene, after DSB induction and repair. This difference in dynamics also appears to be under the control of C(2)M. Indeed, overexpressing C(2)M in nurse cells or the mitotic region of the germline promotes localization of C(3)G and some SC assembly. Thus, C(2)M modifies a complex that is otherwise composed of mitotic cohesins that may control when and where SC assembly is initiated.

## MATERIALS AND METHODS

### Drosophila strains and genetics UAS/Gal4 System to Express RNAi

*Drosophila* stocks and crosses were maintained on standard medium at 25°C. Most rescue experiments done in a *c(2)M^Z0810^ / c(2)M^EP2115^* background. (Manheim and Mckim 2003). Germline RNAi was performed using the following stocks from the Transgenic RNAi Project (TRiP) at Harvard Medical School (Ni *et al*. 2011): *Nipped-B* (GL00574)*, Stromalin* (GL00534)*, vtd* (GL00522) and *SMC3* (GL00518). These transgenic lines express shRNA against the desired RNA and are under control of the GAL4/UAS system. For germline expression, the following GAL4 lines were used: *P{w^+mC^=GAL4::VP16-nos.UTR}CG6325^MVD1^* (Rorth 1998), *P{w[+mC]=GAL4-nos.NGT}A* (Wheeler *et al*. 2002), *P{w^+mc^,GAL4-Hsp70,PB}89-2-1* (Brand and Perrimon 1993), and *P{w^+^, bam:Gal4-VP16}* (Mathieu et al. 2013). For ubiquitous expression we used *P{w^+mC^=tubP-GAL4}LL7* (Lee and Luo 1999).

### Generation of c(2)M Mutants and SA-MYC transgene

Site-directed mutagenesis was used to create *c(2)M* mutants. The template was a wild-type *c(2)M* sequence cloned into the pENTR4 vector. The Q5 Site-Directed Mutagenesis Kit, from New England Biolabs was used with primers created to amplify the *c(2)M* sequence with the desired mutation. Construction of *SMC1* transgenes was described previously (Schuldiner *et al*. 2008; Gyuricza *et al*. 2016), except for these experiments a myc-tag version of SMC1 was used by cloning into the pPWM vector (DGRC).

Full-length cDNA of SA was obtained from the Drosophila Gene Collection (LD34181). The sequence was amplified and with primers to introduce a Xho1 site after the coding region and abolish the stop codon, and blunt cloned into PST BLUE. Using the restriction sites Eco1 Xho1, a C-terminal fragment was subcloned into pENTR4 (Invitrogen). The N-terminal half was excised from the cDNA clone using the restriction sites Bgl1 and HindI and also ligated into the same pENTR4 construct. An expression vector encoding full-length SA fused to a C-terminal MYC tag under the control of the UASp promoter was created by a Clonase LR reaction with the pPWM vector (DGRC). Germline transformations were performed by Model Systems Genomics (Durham, NC) to generate transgene lines.

### Germarium Preparation for Immunofluorescence

In order to visualize and image the loading of C(2)M, SMC, and SA proteins and their localization, germarium stage oocytes were prepared for cytology by the following process. On day 12 of the experimental cross, female virgin and young flies were collected and heat shocked for 1 hour at 37°C, then returned to 25°C following heat shock. 24 hours after completion of heat shock, approximately 20 females were dissected for their oocytes in 1X Robbs buffer for no longer than 20 minutes. Each experimental group was done separately. The oocytes are then fixed in a fixation buffer containing 37% formaldehyde for 9 minutes before getting washed in BAT and BAT-NGS twice in each for 15 minutes. w^-^ embryos were rehydrated in 0.1% tween20 and blocked for 60 minutes in BAT-NGS. The primary antibodies are added to the oocytes in 300 µL of BAT-NGS and then incubated at 4°C overnight while nutating. The primary antibodies for these preps were rat HA, mouse myc, guinea pig CENP-C 703, mouse C(3)G, rabbit C(2)M^910^, and were used according to the sample. The secondary antibodies were added and pre-absorbed by the w^-^ embryos to account for nonspecific binding and incubated at 4°C overnight while nutating in a photosensitive tube. The secondary antibodies for this prep were anti-rat Cy3, anti-mouse Alexa 488, and anti-guinea pig Alexa 647, and anti-rabbit Alexa 647. The following day, the oocytes were washed 4 times in BAT-BSA for 30 minutes each, and then 1 time in BAT-NGS for 30 minutes. Embryos with secondary antibodies were spun down for 5 minutes at 13.9 speed and then added to the oocytes following proper dilution. Oocytes were incubated at room temperature for 4 hours, nutating. After incubation, oocytes were washed 1 time in BAT for 30 minutes. Then, 2.5µL DNA dye Hoecsht was added in 1mL BAT for 7 minutes. Oocytes were washed 1 time in BAT for 15 minutes, then quick-washed in 1X Buffer-A. Oocytes were stored in 1X Buffer-A at 4°C in the dark until ready for mounting to image.

### Imaging the Germarium

Oocytes were mounted on a glass slide in SlowFade (Invitrogen) Refractive Index: 1.42. Individual ovarioles were separated in order to avoid overlapping germariums during imaging. Images were captured with the Leica TCS SP8 and Leica Stellaris confocal microscopes with a 63X, N.A. 1.4 lens and captured full germariums including X, Y, and Z planes. Data on colocalization of SA and SMC staining with C(2)M were quantified based on individual cell staining overlap between antibody channels. Images were later analyzed using Leica software.

These methods have been found to be functional in previous experiments within the McKim Laboratory in germarium oocyte preparation as well as the use of the UAS-Gal4 system. Results from experiments and successful mutant *Drosophila* generation, using the same imaging and analysis tools, have been published (Gottlieb and Tegay 2018).

### Nondisjunction and crossover assays

The frequency of X chromosome nondisjunction [69] and female fertility in flies of certain genotypes was determined by crossing female virgin flies to *yw/B^S^Y* males. If nondisjunction occurred, females would be observed with bar-eyes (XXY), and males with normal non bar-eyes. The XXX and OY genotypes are lethal, which is accounted for in this equation to calculate nondisjunction:

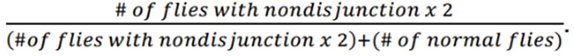

The flies were counted 18 days after each respective cross. To examine the effects on fertility, the number of total flies produced per cross was divided by the original number of females in the parental nondisjunction cross counted on day 18. Nondisjunction rates in mutants were compared to that of wild-type. For measuring corssing over and meiosis II nondisjunction, the X-chromosome genotype was *y w cv m f • y^+^/ y w.* y females crossed to *y Hw w/B^Y^* males. The *y^+^* marker is to the right of the X-chromosome centromere. The yellow progeny result from sister chromatid nondisjunction.

## Acknowledgements

We thank Marina Druzhinina for technical assistance. Stocks obtained from the Bloomington Drosophila Stock Center (NIH P40OD018537) were used in this study. Image acquisition and analysis was made possible by the Waksman Institute Shared Imaging Core Facility and The Human Genetics Institute Imaging Core Facility at Rutgers. This work was supported by NIH grant GM101955 to K.S.M.

## Works Cited

Anderson, L. K., S. M. Royer, S. L. Page, K. S. McKim, A. Lai et al., 2005 Juxtaposition of C(2)M and the transverse filament protein C(3)G within the central region of Drosophila synaptonemal complex. Proc Natl Acad Sci U S A 102: 4482–4487.

Arumugam, P., S. Gruber, K. Tanaka, C. H. Haering, K. Mechtler et al., 2003 ATP hydrolysis is required for cohesin’s association with chromosomes. Curr Biol 13: 1941–1953.

Arumugam, P., T. Nishino, C. H. Haering, S. Gruber and K. Nasmyth, 2006 Cohesin’s ATPase activity is stimulated by the C-terminal Winged-Helix domain of its kleisin subunit. Curr Biol 16: 1998–2008.

Bayes, M., I. Prieto, J. Noguchi, J. L. Barbero and L. A. Perez Jurado, 2001 Evaluation of the Stag3 gene and the synaptonemal complex in a rat model (as/as) for male infertility. Mol Reprod Dev 60: 414–417.

Brand, A. H., and N. Perrimon, 1993 Targeted gene expression as a means of altering cell fates and generating dominant phenotypes. Development 118: 401–415.

Cahoon, C. K., and R. S. Hawley, 2016a Meiosis: Cohesins Are Not Just for Sisters Any More. Curr Biol 26: R523–r525.

Cahoon, C. K., and R. S. Hawley, 2016b Regulating the construction and demolition of the synaptonemal complex. Nat Struct Mol Biol 23: 369–377.

Castellano-Pozo, M., G. Sioutas, C. Barroso, J. P. Prince, P. Lopez-Jimenez et al., 2023 The kleisin subunit controls the function of C. elegans meiotic cohesins by determining the mode of DNA binding and differential regulation by SCC-2 and WAPL-1. Elife 12.

Davis, L., A. E. Rozalén, S. Moreno, G. R. Smith and C. Martín-Castellanos, 2008 Rec25 and Rec27, novel linear-element components, link cohesin to meiotic DNA breakage and recombination. Curr Biol 18: 849–854.

Eijpe, M., H. Offenberg, R. Jessberger, E. Revenkova and C. Heyting, 2003 Meiotic cohesin REC8 marks the axial elements of rat synaptonemal complexes before cohesins SMC1beta and SMC3. J Cell Biol 160: 657–670.

Gligoris, T. G., J. C. Scheinost, F. Bürmann, N. Petela, K. L. Chan et al., 2014 Closing the cohesin ring: structure and function of its Smc3-kleisin interface. Science 346: 963–967.

Gottlieb, S. F., and D. H. Tegay, 2018 Genetics, Nondisjunction in *StatPearls*, Treasure Island (FL).

Gutiérrez-Caballero, C., Y. Herrán, M. Sánchez-Martín, J. A. Suja, J. L. Barbero et al., 2011 Identification and molecular characterization of the mammalian α-kleisin RAD21L. Cell Cycle 10: 1477–1487.

Gyuricza, M. R., K. B. Manheimer, V. Apte, B. Krishnan, E. F. Joyce et al., 2016 Dynamic and Stable Cohesins Regulate Synaptonemal Complex Assembly and Chromosome Segregation. Curr Biol 26: 1688–1698.

Haering, C. H., D. Schoffnegger, T. Nishino, W. Helmhart, K. Nasmyth et al., 2004 Structure and stability of cohesin’s Smc1-kleisin interaction. Mol Cell 15: 951–964.

Heidmann, D., S. Horn, S. Heidmann, A. Schleiffer, K. Nasmyth et al., 2004 The Drosophila meiotic kleisin C(2)M functions before the meiotic divisions. Chromosoma 113: 177–187.

Higashi, T. L., P. Eickhoff, J. S. Sousa, J. Locke, A. Nans et al., 2020 A Structure-Based Mechanism for DNA Entry into the Cohesin Ring. Mol Cell 79: 917–933.e919.

Higashi, T. L., and F. Uhlmann, 2022 SMC complexes: Lifting the lid on loop extrusion. Current Opinion in Cell Biology 74: 13–22.

Hughes, S. E., D. E. Miller, A. L. Miller and R. S. Hawley, 2018 Female Meiosis: Synapsis, Recombination, and Segregation in Drosophila melanogaster. Genetics 208: 875–908.

Ishiguro, K., J. Kim, S. Fujiyama-Nakamura, S. Kato and Y. Watanabe, 2011 A new meiosis-specific cohesin complex implicated in the cohesin code for homologous pairing. EMBO Rep 12: 267–275.

Ishiguro, K., J. Kim, H. Shibuya, A. Hernández-Hernández, A. Suzuki et al., 2014 Meiosis-specific cohesin mediates homolog recognition in mouse spermatocytes. Genes Dev 28: 594–607.

Ishiguro, K. I., 2019 The cohesin complex in mammalian meiosis. Genes Cells 24: 6–30.

Ito, M., and A. Shinohara, 2022 Chromosome architecture and homologous recombination in meiosis. Front Cell Dev Biol 10: 1097446.

Krishnan, B., S. E. Thomas, R. Yan, H. Yamada, I. B. Zhulin et al., 2014 Sisters Unbound Is Required for Meiotic Centromeric Cohesion in Drosophila melanogaster. Genetics.

Lake, C. M., and R. S. Hawley, 2012 The molecular control of meiotic chromosomal behavior: events in early meiotic prophase in Drosophila oocytes. Annu Rev Physiol 74: 425–451.

Lee, J., and T. Hirano, 2011 RAD21L, a novel cohesin subunit implicated in linking homologous chromosomes in mammalian meiosis. J Cell Biol 192: 263–276.

Lee, J., T. Iwai, T. Yokota and M. Yamashita, 2003 Temporally and spatially selective loss of Rec8 protein from meiotic chromosomes during mammalian meiosis. J Cell Sci 116: 2781–2790.

Lee, T., and L. Luo, 1999 Mosaic analysis with a repressible cell marker for studies of gene function in neuronal morphogenesis. Neuron 22: 451–461.

Li, Y., K. W. Muir, M. W. Bowler, J. Metz, C. H. Haering et al., 2018 Structural basis for Scc3-dependent cohesin recruitment to chromatin. Elife 7.

Losada, A., M. Hirano and T. Hirano, 1998 Identification of Xenopus SMC protein complexes required for sister chromatid cohesion. Genes Dev 12: 1986–1997.

Losada, A., and T. Hirano, 2005 Dynamic molecular linkers of the genome: the first decade of SMC proteins. Genes Dev 19: 1269–1287.

Manheim, E. A., and K. S. McKim, 2003 The Synaptonemal Complex Component C(2)M Regulates Meiotic Crossing over in Drosophila. Curr Biol 13: 276–285.

Mathieu, J., C. Cauvin, C. Moch, S. J. Radford, P. Sampaio et al., 2013 Aurora B and cyclin B have opposite effects on the timing of cytokinesis abscission in Drosophila germ cells and in vertebrate somatic cells. Dev Cell 26: 250–265.

McKearin, D. M., and A. C. Spradling, 1990 bag-of-marbles; a Drosophila gene required to initiate male and female gametogenesis. Genes & Dev. 4: 2242–2251.

Ni, J. Q., R. Zhou, B. Czech, L. P. Liu, L. Holderbaum et al., 2011 A genome-scale shRNA resource for transgenic RNAi in Drosophila. Nat Methods 8: 405–407.

Ohlstein, B., and D. McKearin, 1997 Ectopic expression of the Drosophila Bam protein eliminates oogenic germline stem cells. Development 124: 3651–3662.

Parisi, S., M. J. McKay, M. Molnar, M. A. Thompson, P. J. van der Spek et al., 1999 Rec8p, a meiotic recombination and sister chromatid cohesion phosphoprotein of the Rad21p family conserved from fission yeast to humans. Mol Cell Biol 19: 3515–3528.

Prieto, I., J. A. Suja, N. Pezzi, L. Kremer, A. C. Martinez et al., 2001 Mammalian STAG3 is a cohesin specific to sister chromatid arms in meiosis I. Nat Cell Biol 3: 761–766.

Revenkova, E., M. Eijpe, C. Heyting, B. Gross and R. Jessberger, 2001 Novel meiosis-specific isoform of mammalian SMC1. Mol Cell Biol 21: 6984–6998.

Rorth, P., 1998 Gal4 in the Drosophila female germline. Mech Dev 78: 113–118.

Schleiffer, A., S. Kaitna, S. Maurer-Stroh, M. Glotzer, K. Nasmyth et al., 2003 Kleisins: a superfamily of bacterial and eukaryotic SMC protein partners. Mol Cell 11: 571–575.

Schuldiner, O., D. Berdnik, J. M. Levy, J. S. Wu, D. Luginbuhl et al., 2008 piggyBac-based mosaic screen identifies a postmitotic function for cohesin in regulating developmental axon pruning. Dev Cell 14: 227–238.

Tanneti, N. S., K. Landy, E. F. Joyce and K. S. McKim, 2011 A Pathway for Synapsis Initiation during Zygotene in Drosophila Oocytes. Curr Biol 21: 1852–1857.

Ur, S. N., and K. D. Corbett, 2021 Architecture and Dynamics of Meiotic Chromosomes. Annu Rev Genet 55: 497–526.

Webber, H. A., L. Howard and S. E. Bickel, 2004 The cohesion protein ORD is required for homologue bias during meiotic recombination. J Cell Biol 164: 819–829.

Wells, J. N., T. G. Gligoris, K. A. Nasmyth and J. A. Marsh, 2017 Evolution of condensin and cohesin complexes driven by replacement of Kite by Hawk proteins. Curr Biol 27: R17–R18.

Weng, K. A., C. A. Jeffreys and S. E. Bickel, 2014 Rejuvenation of Meiotic Cohesion in Oocytes during Prophase I Is Required for Chiasma Maintenance and Accurate Chromosome Segregation. PLoS Genet 10: e1004607.

Wheeler, J. C., C. VanderZwan, X. Xu, D. Swantek, W. D. Tracey et al., 2002 Distinct in vivo requirements for establishment versus maintenance of transcriptional repression. Nat Genet 32: 206–210.

Winters, T., F. McNicoll and R. Jessberger, 2014 Meiotic cohesin STAG3 is required for chromosome axis formation and sister chromatid cohesion. EMBO J 33: 1256–1270.

Wintrebert, M., M. C. Nguyen and G. R. Smith, 2021 Activation of meiotic recombination by nuclear import of the DNA break hotspot-determining complex in fission yeast. J Cell Sci 134.

Yan, R., and B. D. McKee, 2013 The Cohesion Protein SOLO Associates with SMC1 and Is Required for Synapsis, Recombination, Homolog Bias and Cohesion and Pairing of Centromeres in Drosophila Meiosis. PLoS Genet 9: e1003637.

Zhao, Y., L. Ren, T. Zhao, H. You, Y. Miao et al., 2024 SCC3 is an axial element essential for homologous chromosome pairing and synapsis. Elife 13.

Zickler, D., and N. Kleckner, 2023 Meiosis: Dances Between Homologs. Annu Rev Genet 57: 1–63.

