## Supplemental Table and Figures for "Exploring the C(2)M Cohesin Complex: Structure, Dynamics, and Ability to Facilitate Assembly of the Synaptonemal Complex"

Table S 1: **C(2)M Mutants in Wild-Type Background**

| | Oocyte genotype | | | | NDJ (%) |
|---|---|---|---|---|---|
| | XXY ♀ | XO ♂ | XX ♀ | XY ♂ | |
| L97K-L528R | 7 | 4 | 1915 | 1769 | 0.59 |
| V86K | 0 | 0 | 697 | 596 | 0 |
| L97K | 3 | 4 | 799 | 589 | 1.00 |
| D100K | 0 | 1 | 309 | 222 | 0.38 |
| F524A | ND | ND | ND | ND | ND |
| F524R | 1 | 4 | 85 | 61 | 6.41 |
| L528A | 2 | 6 | 1552 | 1347 | 0.55 |
| L528R | 0 | 2 | 1692 | 1435 | 0.13 |
| Y528R | 1 | 1 | 892 | 694 | 0.25 |
| C(2)M-ΔN | 9 | 17 | 2153 | 1929 | 1.26 |
| CRC | 3 | 12 | 1931 | 1696 | 0.8 |

Figure S1

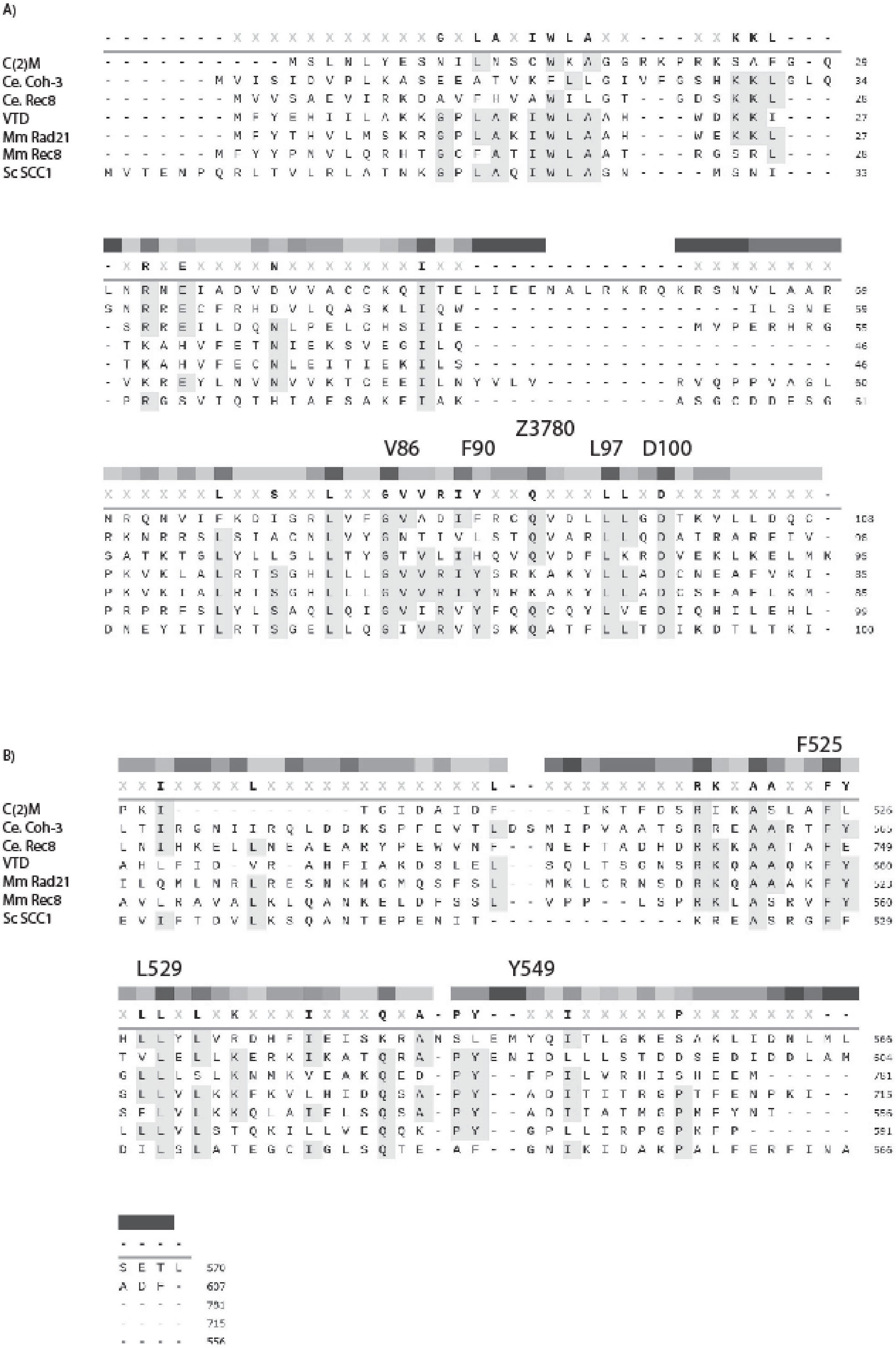
A)
C(2)M
Ce. Coh-3
Ce. Rec8
VTD
Mm Rad21
Mm Rec8
Sc SCC1
V86
F90
Z3780
L97
D100
B)
F525
L529
Y549

Figure S 1: Alignment of C(2)M N- and C-terminal domains.

Alignment of C(2)M with orthogues: *C. elegans* (Coh-3) and mouse Rad21L, and paralogs *C. elegans* Rec-8, D. melanogaster VTD/Rad21, mouse Rad21, budding yeast Scc1. Only alignments of the (A) N- and (B) C- terminal domains are shown because the central domain is poorly conserved.

Figure S2

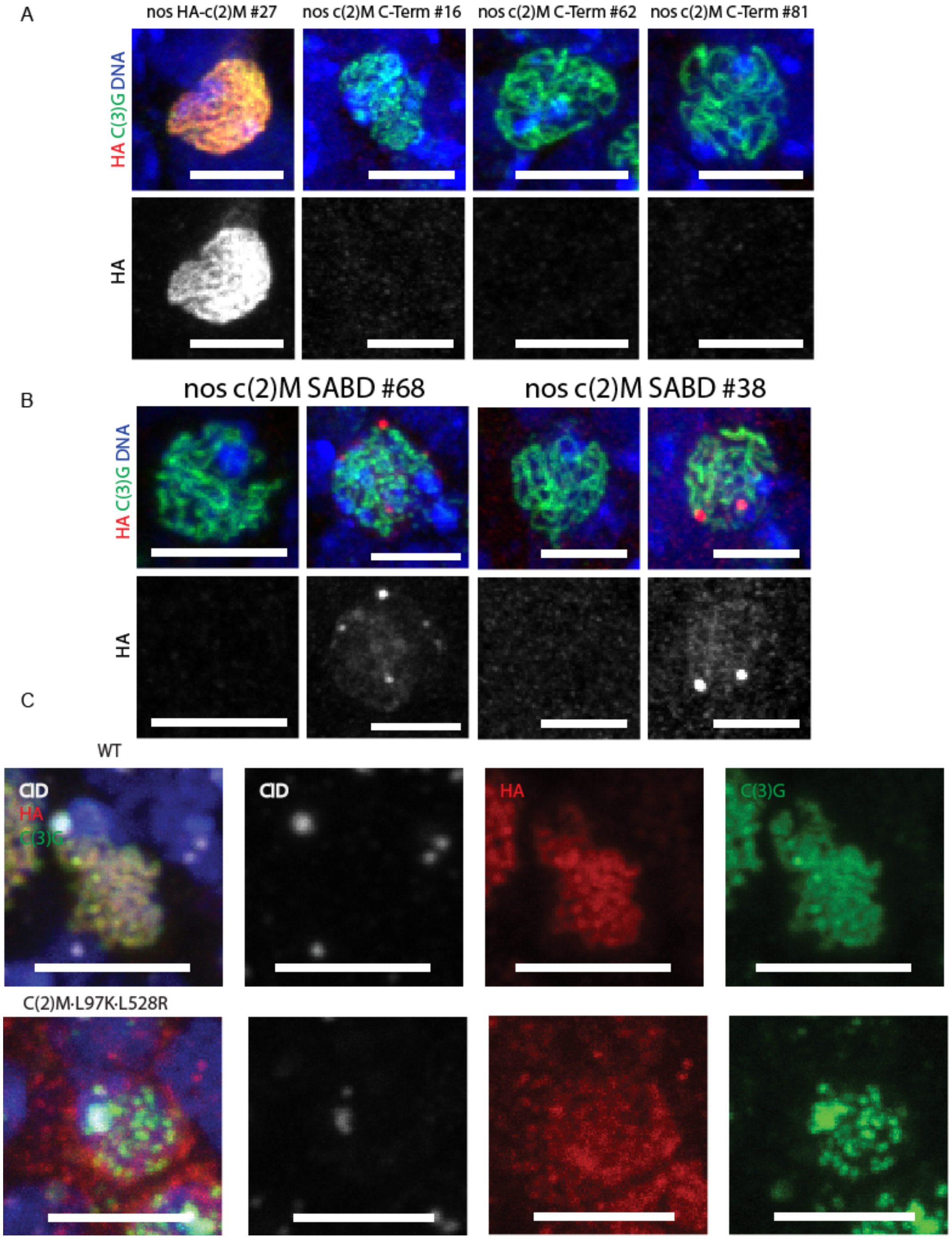


### Figure S 2: Localization of C-terminal or SA-Binding domains of C(2)M in wild-type background.

A) Expression of the C terminus of C(2)M. The C terminus did not express in three independent lines. B) Expression of central SA domain results in formation of  aggregates indicated by the foci in red. C(3)G is in green, HA / C(2)M is in red, DNA in blue, and the scale bar is 5 μM. C) Localization of mutants with missense mutations in the N (L97K) and C-terminal (L529R) domains.

Figure S3

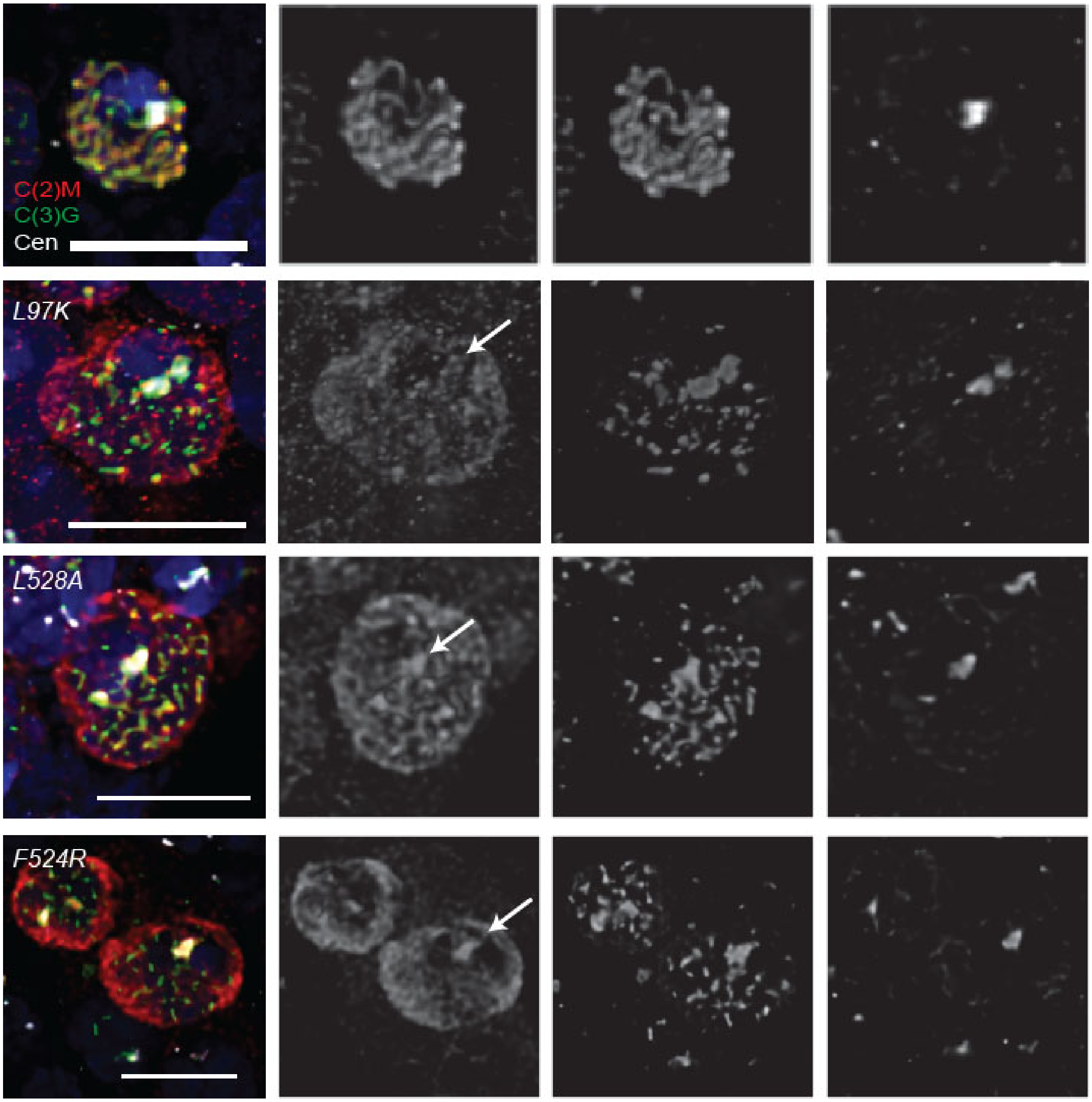

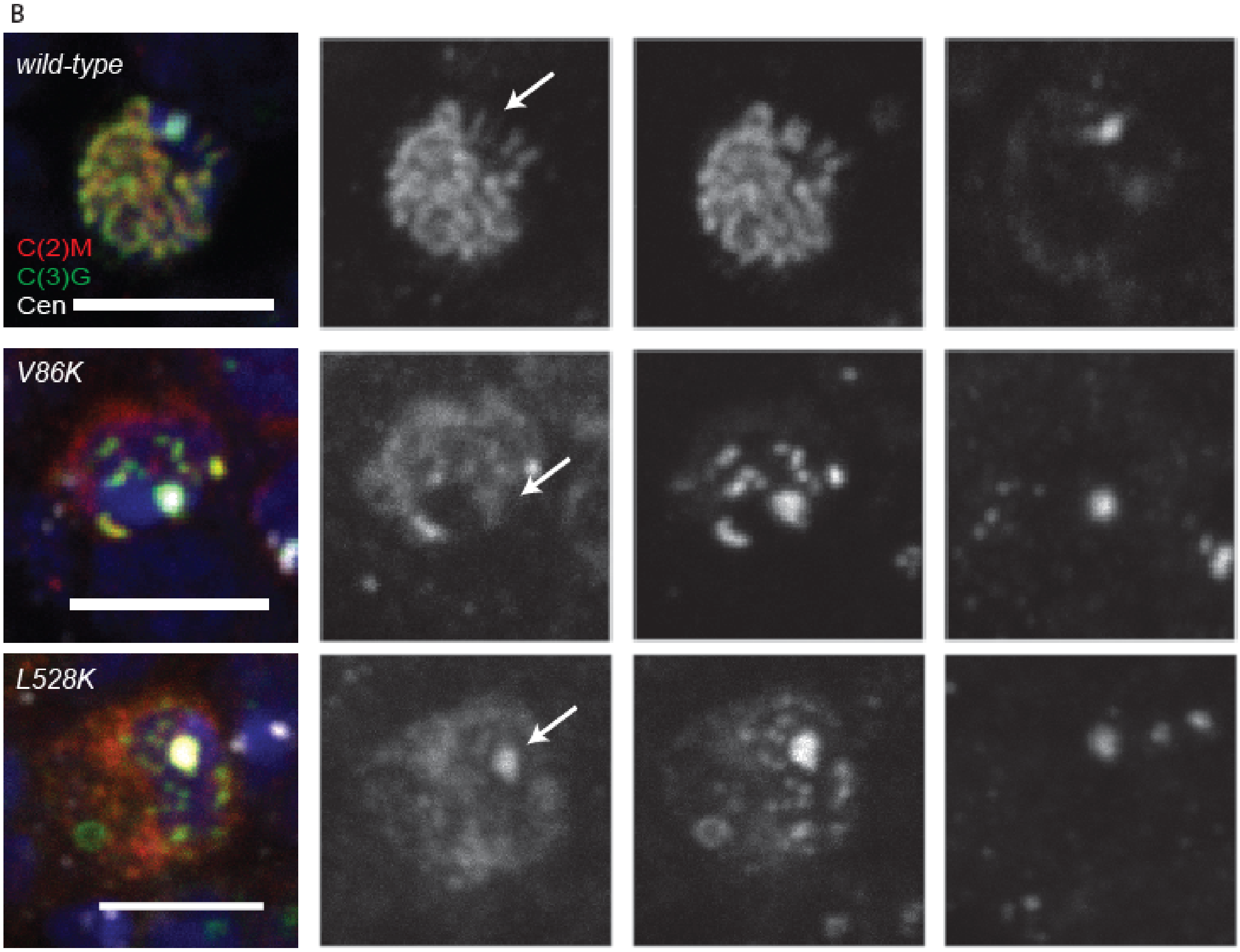


### Figure S 3: Extra images of *c(2)M* mutants with centromere staining.

Images of pachytene nuclei using a limited number of optical sections that include the centromeres (CENP-C in white). Arrow shows the location of the centromere in the C(2)M channel. C(3)G is in green, HA-C(2)M is in red, DNA in blue, and the scale bar is 5 μM.

Figure S4

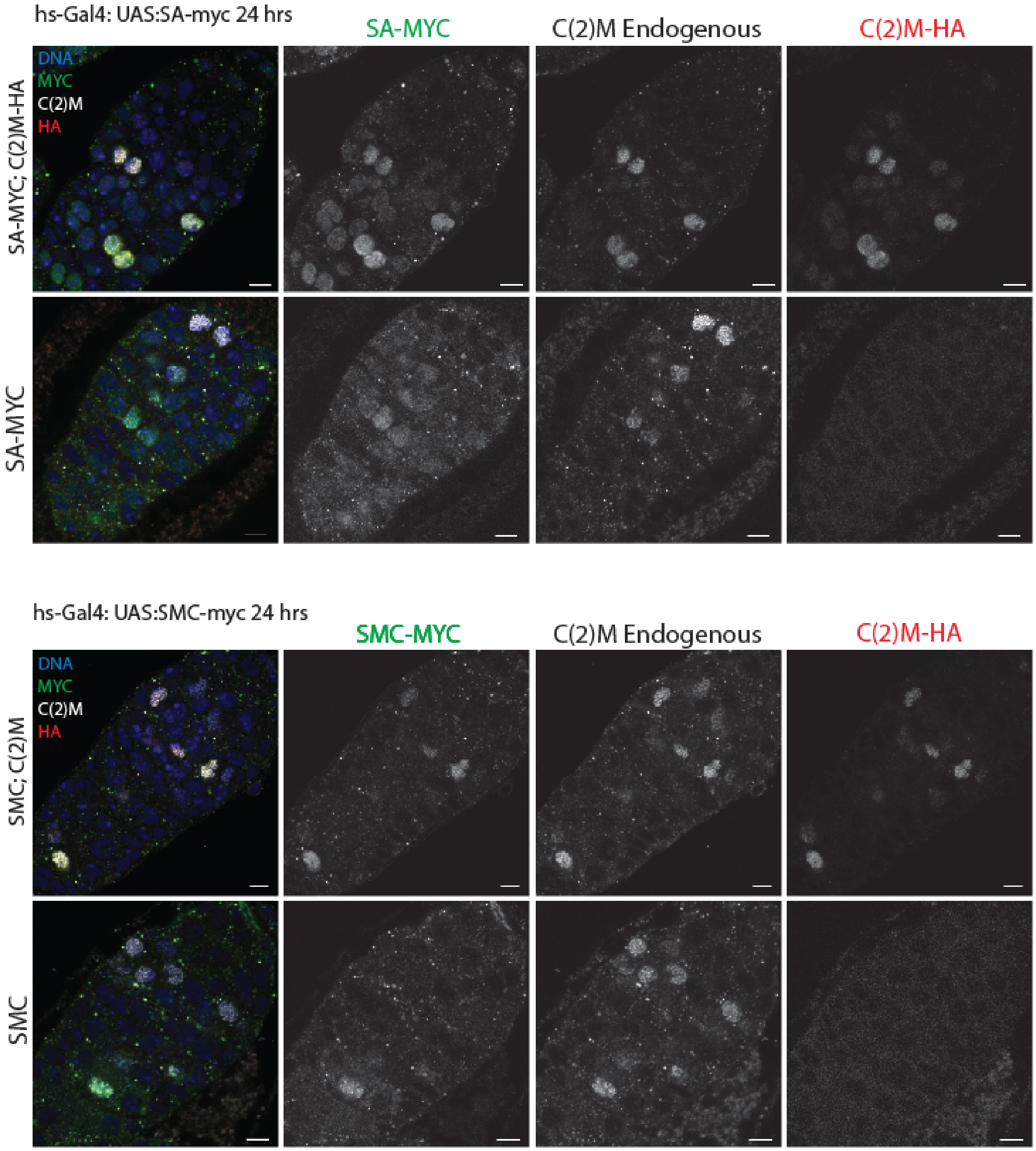


### Figure S 4: Heat shock induced SA and SMC3, whole germarium view.

Same parameters and channels as **Error! Reference source not found.** but showing the whole germarium. C(3)G is in green, HA-C(2)M is in red, DNA in blue, and the scale bar is 5 µM.

Figure S5

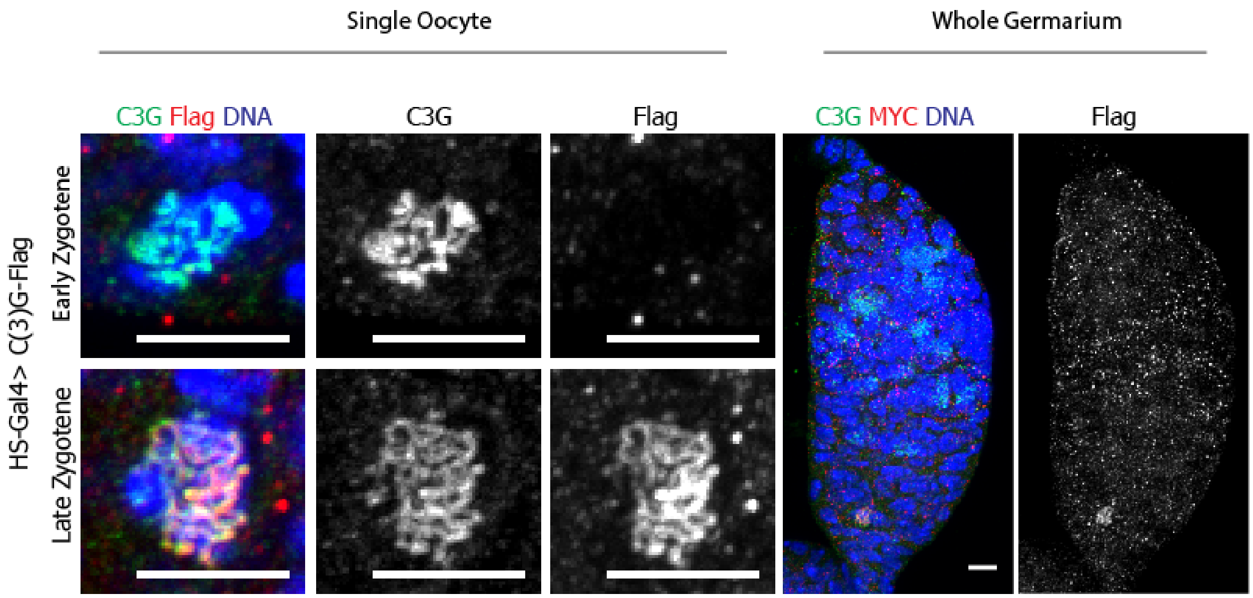


### Figure S 5: Heat shock induced expression of C(3)G

Endogenous C(3)G is in green, while heat shock induced FLAG-C(3)G is in red. DNA is in blue is the DNA and the scale bars are 5μm. Shown in each panel is a selected oocyte from an early stage (2A) and a later stage (3) as well as a whole germarium.
